# Stratified analysis identifies HIF-2*α* as a therapeutic target for highly immune-infiltrated melanomas

**DOI:** 10.1101/2024.10.29.620300

**Authors:** Amy Y. Huang, Kelly P. Burke, Ryan Porter, Lynn Meiger, Peter Fatouros, Jiekun Yang, Emily Robitschek, Natalie Vokes, Cora Ricker, Valeria Rosado, Giuseppe Tarantino, Jiajia Chen, Tyler J. Aprati, Marc C. Glettig, Yiwen He, Cassia Wang, Doris Fu, Li-Lun Ho, Kyriakitsa Galani, Gordon J. Freeman, Elizabeth I. Buchbinder, F. Stephen Hodi, Manolis Kellis, Genevieve M. Boland, Arlene H. Sharpe, David Liu

## Abstract

While immune-checkpoint blockade (ICB) has revolutionized treatment of metastatic melanoma over the last decade, the identification of broadly applicable robust biomarkers has been challenging, driven in large part by the heterogeneity of ICB regimens and patient and tumor characteristics. To disentangle these features, we performed a standardized meta-analysis of eight cohorts of patients treated with anti-PD-1 (n=290), anti-CTLA-4 (n=175), and combination anti-PD-1/anti-CTLA-4 (n=51) with RNA sequencing of pre-treatment tumor and clinical annotations. Stratifying by immune-high vs -low tumors, we found that surprisingly, high immune infiltrate was a biomarker for response to combination ICB, but not anti-PD-1 alone. Additionally, hypoxia-related signatures were associated with non-response to anti-PD-1, but only amongst immune infiltrate-high melanomas. In a cohort of scRNA-seq of patients with metastatic melanoma, hypoxia also correlated with immunosuppression and changes in tumor-stromal communication in the tumor microenvironment (TME). Clinically actionable targets of hypoxia signaling were also uniquely expressed across different cell types. We focused on one such target, HIF-2*α*, which was specifically upregulated in endothelial cells and fibroblasts but not in immune cells or tumor cells. HIF-2*α* inhibition, in combination with anti-PD-1, enhanced tumor growth control in pre-clinical models, but only in a more immune-infiltrated melanoma model. Our work demonstrates how careful stratification by clinical and molecular characteristics can be leveraged to derive meaningful biological insights and lead to the rational discovery of novel clinical targets for combination therapy.

## Introduction

Immune checkpoint blockade has dramatically transformed the treatment approach for advanced melanoma, leading to impressive and long-lasting treatment outcomes in a subset of patients [1, 2, 3]. Several ICB regimens are approved for treating patients with metastatic melanoma, including anti-CTLA-4 blockade, anti-PD-1 blockade, and combination therapies including anti-CTLA-4/anti-PD-1 blockade and, more recently anti-PD-1/anti-LAG3. ICB regimens in melanoma have important differences in response rate and toxicity: the highest response and survival rates are seen in combination anti-PD-1/anti-CTLA-4 (52% overall survival at 5 years), but so are the highest rates of toxicity (59% of Grade 3 or 4 treatment-related adverse events, compared to 21% and 28% in anti-PD-1 and anti-CTLA-4 monotherapies respectively) [4]. Understanding how underlying patient-to-patient variability contributes to why patients do or do not respond to different checkpoint blockade regimens is important both for clinical practice, in matching patients to the appropriate therapy based on likely response and toxicity, and in the development of more efficacious combination therapies. Identifying features associated with response to ICB has been a topic of great interest over the past decade [5], and many response-associated biological features have been identified through analyzing genomic and transcriptomic data from clinical cohorts. These include tumor mutational burden (TMB) and neoantigen load [6, 7], protein-level expression of PD-L1 and CD8 [8], transcriptional signatures related to mesenchymal transition and extracellular matrix remodeling (IPRES) [9], genetic alternations resulting in loss of antigen presentation [10, 11], interferon gamma (IFN*γ*) signaling pathways [12, 13], tumor heterogeneity and ploidy [14], and, recently, deactivation of ligand-receptor interactions enhancing lymphocyte infiltration [15]. However, these observations have often been made in heterogeneous clinical cohorts, combining different melanoma subtypes, ICB regimens, and prior treatments. More recent studies have found that the association of these signatures with response is impacted by clinical treatment history, highlighting differences in responders to anti-PD-1 monotherapy and combination ICB [16, 17] as well as differences in patients with and without previous exposure to anti-CTLA-4 blockade [18, 19]. Additional work has focused on patient characteristics, such as age [20].

Given the growing amount of clinical data with high-dimensional molecular characterization, we hypothesized that stratified meta-analysis of a well-annotated cohort of patients would enable us to disentangle features predicting differential response to ICB in heterogeneous populations of patients. Thus, we performed an integrative analysis seeking to understand the biology of response to ICB under different treatment regimens (Fig. 1a). First, we combined sequencing and clinical data from eight previously published cohorts (693 patient samples total) [6, 21, 22, 16, 18, 23, 19, 24], with comprehensively annotated and sequenced pre-treatment tumors from patients with metastatic melanoma treated with anti-CTLA-4, anti-PD-1, and combination ICB. Upon observing large variation in immune infiltrate across the cohort through bulk RNA-sequencing analysis, we then stratified the cohort into “immune-high” and “immune-low,” and characterized transcriptomic features to identify biological pathways associated with response and resistance within those groups. We then utilized several unrelated cohorts of metastatic melanoma patients (55 samples total) with single cell RNA-sequencing of pre-treatment tumors to further investigate these pathways. Among other key findings, our analysis suggested that hypoxia was a biomarker of non-response to anti-PD-1 specifically in patients with immune-high tumors and correlated with functional changes in the TME. We identified HIF-2*α* as a novel hypoxia-related target uniquely expressed by fibroblasts and endothelial cells but not by tumor or immune cells in melanomas. We then tested the potential for targeting HIF-2*α* combined with anti-PD-1 in pre-clinical models. Combination treatment delayed tumor growth in an immune-high but not an immune-low model.

**Figure 1.**
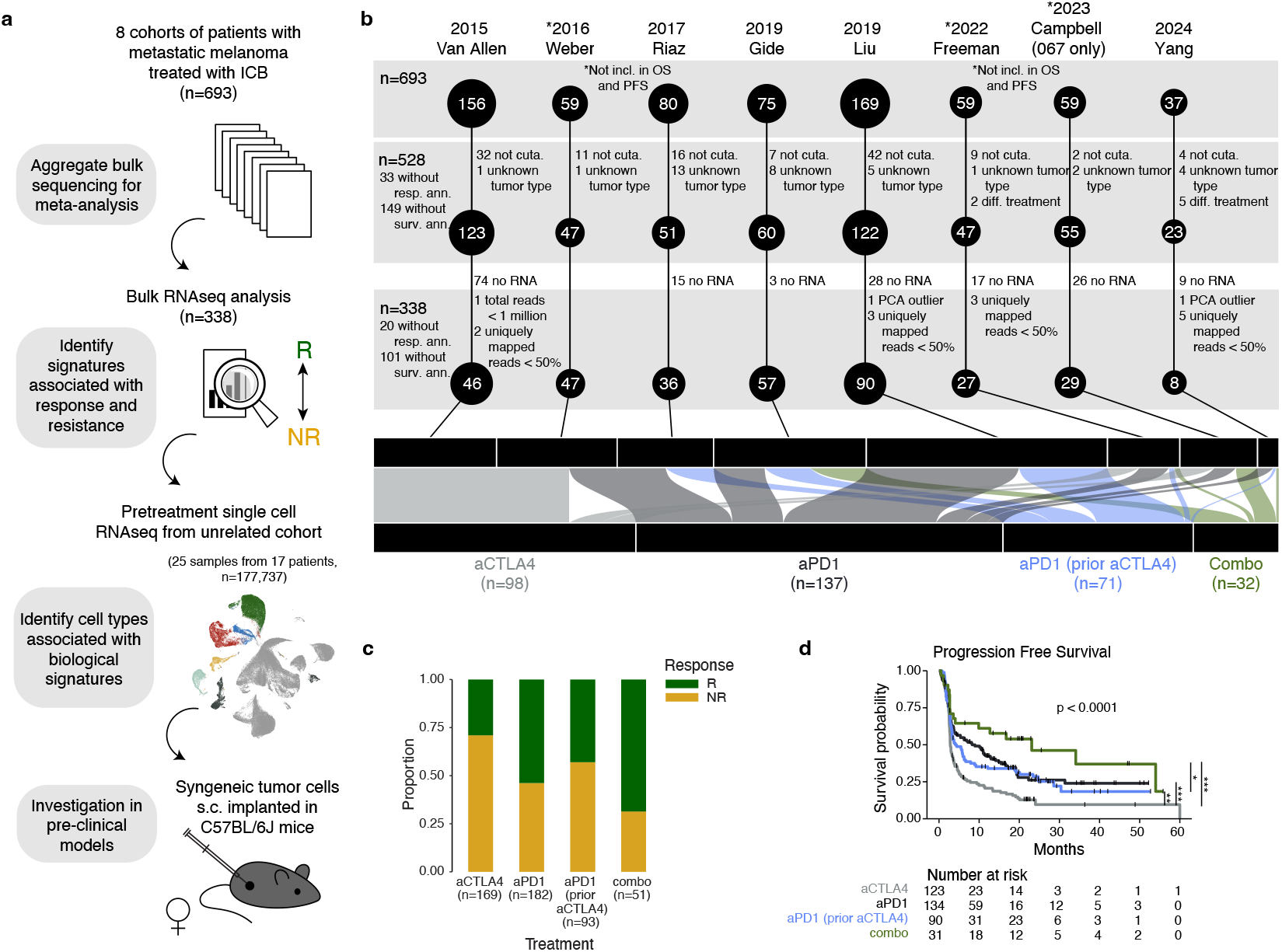
Study overview, cohort selection, and clinical characteristics. **a**. Diagram depicting study overview. The study has four major components: (1) aggregate bulk RNA-sequencing data from 8 cohorts for a meta-analysis, (2) identify features of the tumor microenvironment (TME) associated with response and resistance, (3) analyze cell types associated with such features in orthogonal scRNA-sequencing cohorts to evaluate the potential for targeting such signatures, and (4) testing in pre-clinical models. **b**. Eight cohorts included in bulk transcriptomic analysis (n=694). **c**. Response rates by treatment group. Only samples which passed clinical inclusion criteria with response data were included; 33 samples were missing response annotations. **d**. Progression-free survival rates stratified by treatment group. Samples which passed clinical exclusion criteria with survival data were considered. 149 samples were missing survival data or were treated sequentially; these were excluded from the analysis. The survival rates between the four treatment groups were significantly different (Log-rank *p <* 0.0001). Significance of pairwise log-rank test also shown, where * * \**p <* 0.001, * * *p <* 0.01, \**p <* 0.05.

## Results

### Clinical characteristics of aggregated cohort

We formed a meta-cohort from eight cohorts of immune checkpoint blockade (ICB)-treated (anti-PD-1 alone, anti-CTLA-4 alone, and combination) melanoma patients with whole-transcriptome sequencing (RNA-seq) or whole-exome sequencing (WES) of pre-treatment tumor and clinical annotations (n=693) [6, 21, 22, 16, 18, 23, 19, 24]. To our knowledge, these eight cohorts comprise all available cohorts of pre-treatment tumor with sequencing and comprehensive clinical annotations. Overall, 77.1% were cutaneous melanomas (534/693), 5.9% were mucosal (41/693), 3.2% were acral (22/693), 3.8% were uveal (26/693), and 10.1% were occult or unknown in origin (70/693). For our analysis, we focused on patients with cutaneous melanoma and excluded a small number of patients with prior anti-PD-1 or prior combination anti-CTLA-4/anti-PD-1, as well as patients treated with combination targeted therapy and ICB, leaving a final cohort of n=528 (Fig. 1b). Given that patients with a prior treatment history of anti-CTLA-4 have been shown to have distinct features of response [18], we divided the cohort into four treatment groups: anti-PD-1, anti-CTLA-4, anti-PD-1 with prior anti-CTLA-4, and combination anti-PD-1/anti-CTLA-4 (Fig. 1b). Full details of the meta-cohort can be found in Supplementary Table 1.

495 out of 528 samples were annotated as responders or non-responders (n=222 responders, 273 non-responders, Supplementary Fig. 1a, Methods). As expected, we saw the highest response rates in combination anti-PD-1/anti-CTLA-4 (35/51, 68.6%), followed by single agent anti-PD-1 (98/182, 53.8%) and antiCTLA-4 (49/169, 29.0%), with anti-PD-1 with prior antiCTLA-4 having response rates in between anti-PD-1 and anti-CTLA-4 (40/93, 43.0%) (Fig. 1c), with similar trends within each independent cohort (Supplementary Fig. 1b). Consistent with response rates, overall and progression-free survival (OS and PFS) were significantly different between the four treatment groups in the discovery cohort (Fig. 1d, Supplementary Fig. 1c). Overall, ICB response and survival in our cohort are consistent with previous reports [4].

### Stratification of bulk cohort by immune score

Previous analyses utilizing clinical data have explored genomic features associated with response in melanoma [18, 14, 23]. In our analysis, we stratified by transcriptomic features to evaluate the effects of heterogeneity in the TME on predicting response to ICB. We processed all pre-treatment tumor samples in our meta-cohort with bulk RNA-seq using a standardized computational pipeline with batch correction (Supplementary Fig. 2a-e, Methods). Batch-corrected RNA-seq data with associated metadata can be found in Supplementary Tables 2 and 3. Utilizing cell type and microenvironmental signatures from Bagaev et al. 2021 [25], we calculated a signature score for each sample, for each cell type. In principal component analysis (PCA), the first principal component (PC) explained 50% of the variance of the dataset. The loadings of the first PC indicate the contribution of each variable, with higher absolute values signifying greater importance; we found that immune cell type signatures contributed the most to the first PC (Supplementary Fig. 3a). Immune cell type signatures were also all highly co-correlated (Supplementary Fig. 3b). Through hierarchical clustering solely on the immune signatures, we separated tumors by immune state, categorizing them as either “immune-high” (n=211) or “immune-low” (n=127) (Fig. 2a).

**Figure 2.**
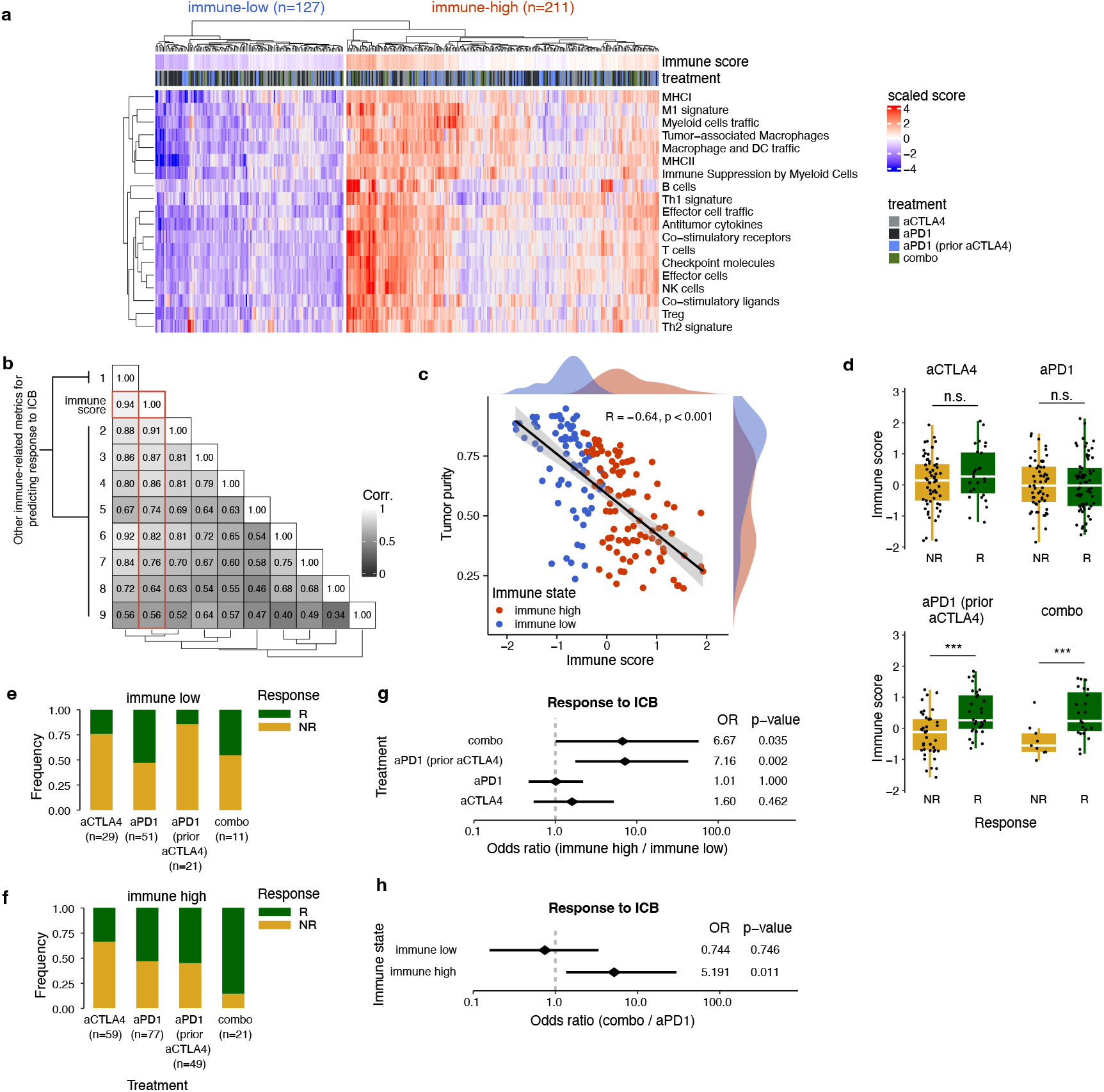
Immune stratification reveals increased response rates to combination ICB in patients with immune-high tumors. **a**. Heatmap showing the scaled ssGSEA scores of 19 immune cell type signatures across all bulk RNA-seq samples. Across all samples, hierarchical clustering is used to define two clusters: immune-high and immune-low. **b**. Co-correlation matrix between the immune score and other immune infiltrate metrics showing high levels of correlation between the immune score and other metrics (between 0.56 and 0.94). From left to right and top to bottom, the columns of the heatmap are: (1) expanded IFN_*γ*_ signature [12], (*) the immune score (outlined in red), (2) Hallmark IFN_*γ*_ signature [26], (3) TLS marker genes [27], (4) CIBERSORTx Tirosh immune proportion [70], (5) CIBERSORTx LM22 absolute immune score [71], (6) non-expanded IFN_*γ*_ signature [12], (7) CIBERSORTx LM22 CD8+ T cell proportion [71], (8) CIBERSORTx Tirosh CD8+ T cell proportion [70], (9) Cabrita TLS signature [27]. **c**. In samples with both whole exome sequencing and bulk RNA-seq (n=162), tumor purity (estimated by FACETS [77]) anti-correlates with the immune score (Spearman’s *ρ* = −0.64, *p <* 0.0001). **d**. The immune score by responders and non-responders in each treatment group. **e-f**. Frequency of response by treatment group in (e) immune-low and (f) immune-high. **g**. Odds ratio and significance by one-sided Fisher’s Exact Test for response in immune-high versus immune-low populations for each treatment group. **h**. Odds ratio and significance by one-sided Fisher’s Exact Test for response in combination versus anti-PD-1 in immune-high and immune-low. For **d-f**., 20 samples with RNA-seq but no response annotations are not shown. * * \**p <* 0.001, * * *p <* 0.01, \**p <* 0.05; n.s. denotes “not significant.”

To represent an overall “immune score”, we calculated an overall global mean of immune signatures (Methods), and we investigated how it related to other metrics which have been proposed to be relevant to predicting response to ICB. Using the same data, we investigated nine other immune-related metrics: the IFN*γ* 10gene and the 28-gene expanded signature from Ayers et al. 2017 [12], the Hallmarks interferon gamma signature [26], tertiary lymphoid structure (TLS) markers genes and a TLS signature from Cabrita et al. 2020 [27], and several signatures estimated using CIBERSORTx [28]: the proportion of immune cells and the proportion of CD8+ T cells inferred using two different reference matrices (Methods). These metrics were highly correlated with each other and the immune score (Fig. 2b). The immune score was most highly correlated with the expanded interferon gamma signature (Spearman’s *ρ* = 0.94) and least correlated with the TLS and the CIBERSORTx-estimated CD8+ T cell proportion (Spearman’s *ρ* = 0.56, 0.64, respectively). Additionally, in samples with paired whole exome sequencing, the immune score was independent of TMB (Pearson’s *R* = 0.027, *p* = 0.75, Supplementary Fig. 3c) but negatively correlated with tumor purity (Pearson’s *R* = − 0.6, *p <* 0.001, Fig. 2c). The immune score also was not associated with biopsy site (Fisher’s Exact Test *p* = 0.735; biopsy site categories including lymph node, skin, and other) or antigen presentation mutations (Fisher’s Exact Test *p* = 0.87; see Methods). The immune score thus provides a coarse-grained representation of the degree of immune infiltrate in the tumor microenvironment.

### An “immune-high” but not “immune-low” score separates response to combination versus single-agent ICB

Next, we examined how the immune score correlated to response (Fig. 2d). Consistent with prior observations [18], the immune score was higher in responders versus non-responders in patients treated with anti-PD-1 with prior anti-CTLA-4 treatment (Wilcoxon rank-sum p=0.00043). However, interestingly, the immune score was not significantly associated with response to single agent anti-PD-1 or anti-CTLA-4 but was strongly associated with response to combination anti-PD-1/anti-CTLA-4 (Wilcoxon rank-sum p=0.00416). To model this effect, we constructed a multi-variable logistic regression model predicting response from the immune score, treatment, and their interaction (Methods, Supplementary Table 4, using anti-PD-1 as a reference. The model predicted that anti-CTLA-4 treatment has worse response rates (OR=0.364, p=0.0008), and that combination ICB has better response rates (OR=3.031, p=0.0528), relative to anti-PD-1, as expected. Furthermore, only in the combination ICB, a greater immune score predicted even better response rates (OR=8.680, p=0.0226). Concordant results were observed stratifying the cohort into “immune-high” and “immune-low” clusters (Fig. 2a). In combination ICB and anti-PD1 with prior anti-CTLA-4, immune-high samples have higher response rates compared to immune-low samples (Fig. 2e-f). For combination blockade, the odds of response were 6.67 higher in patients with immune-high versus immune-low samples (Fig. 2g, Fisher’s Exact Test p=0.035). These data demonstrate that immune infiltrate may be useful for predicting response to combination anti-PD-1/anti-CTLA-4, and affirm that prior history of anti-CTLA-4 treatment alters the response to subsequent anti-PD-1 monotherapy. We also compared response rates between treatment groups (Fig. 2h). Immune-high patients treated with combination ICB had a significantly increased odds of response compared to anti-PD-1 (OR=5.191, Fisher’s Exact Test p=0.011). However, responses between combination ICB and anti-PD-1 were similar in the immune-low group, irrespective of treatment (OR=0.744, Fisher’s Exact Test p=0.746). Therefore, patients with immune-high tumors are much more likely to respond to combination ICB, but those with immune-low tumors are not.

### Hypoxia-related signatures are enriched in immune-high environments of non-responders to anti-PD-1

We reasoned that bulk RNA-sequencing transcriptomes in immune-high and immune-low states likely represent different proportions of immune cells and tumors. No significant differences were seen in individual gene expression between responders and nonresponders for each treatment group after multiple hypothesis test correction (Supplementary Table 5). We thus tested how gene pathways or signatures were associated with response or non-response for each treatment group and whether the addition of our immune score further stratifies response to treatment. Using 254 gene sets of biological processes from the Hallmarks and Kegg databases, we also performed preranked gene set enrichment analysis (GSEA) by treatment groups and immune states. While some signatures were enriched in responders across treatments and immune states (e.g. T cell signaling related signatures, labelled “T cell signaling” on Fig. 3a), other signatures were specific to either immune-low or immune-high states (e.g. a set of metabolic signatures largely enriched in non-responders for immune-low tumors, labelled “metabolism 2” on Fig. 3a). Additional manually identified categories of signatures are shown in Fig. 3a; full results can be found in Supplementary Table 6.

**Figure 3.**
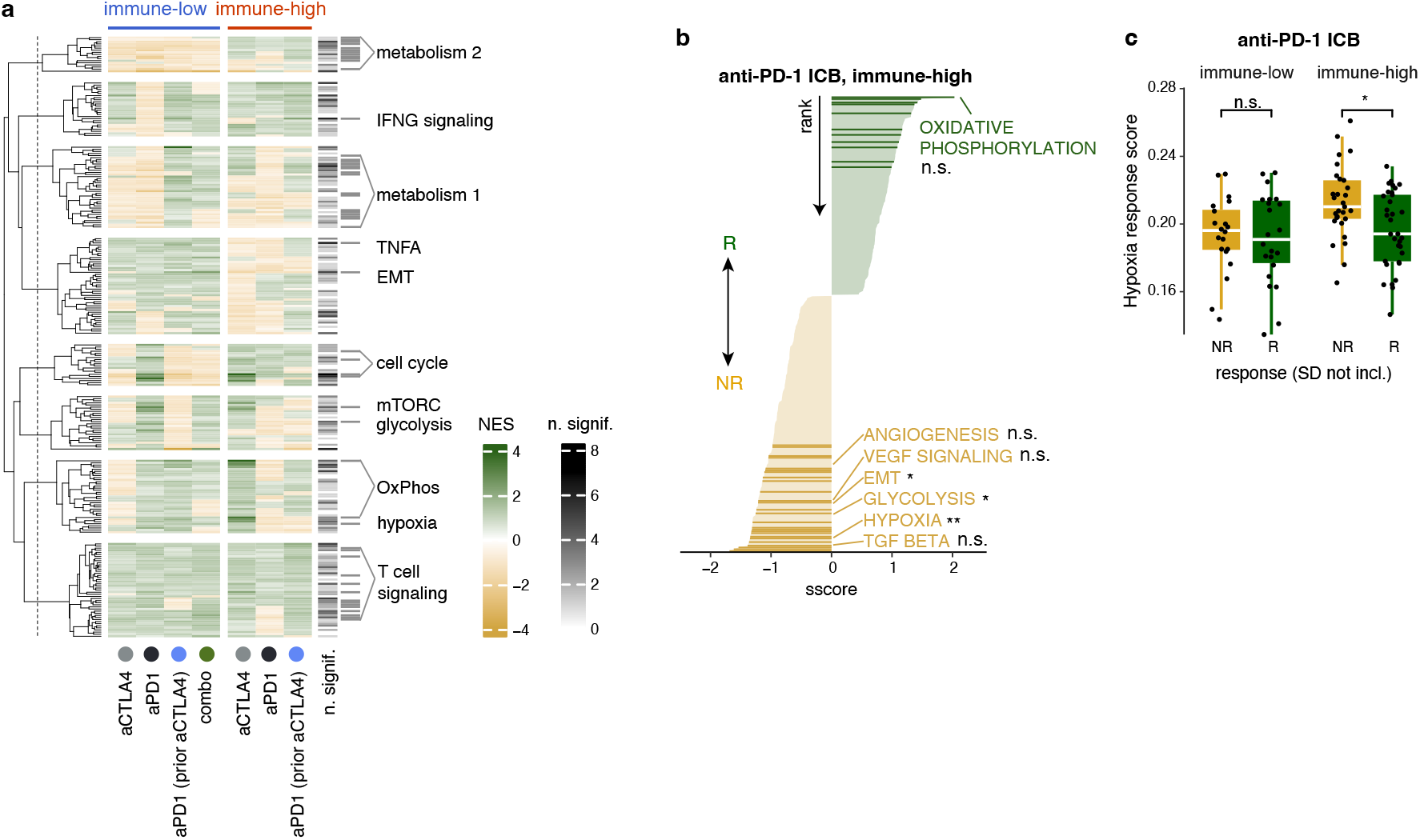
Hypoxia and related pathways enrich in non-responders to anti-PD-1 in immune-high tumor samples. **a**. Enrichment of 254 pathways from the Hallmarks and Kegg databases, including select T cell signatures from ImmuneSigDB. Enrichment analysis was performed for each group of immune-high or low samples for each treatment group separately. Number of significant treatment/immune conditions for which the signature is enriched (0-7) is shown on the annotations on the right of the heatmap. Broad categories for sets of gene signatures of interest are labelled to the right of the significance annotation; grey bars denote signatures which fall into the given categories. **b**. Normalized enrichment score for response to anti-PD-1, immune-high. Hypoxia and related signatures are labelled; *: FDR*<* 0.05, **: FDR*<* 0.01. **c**. ssGSEA score of hypoxia response to anti-PD-1 in immune-high and immune-low. *: *p <* 0.05 by Wilcoxon rank-sum. n.s. denotes “not significant.”

We then examined pathways that were enriched in responders and non-responders differentially between immune-high and immune- low tumors (), with a focus on anti-PD-1 monotherapy (Supplementary Fig. 4a, Supplementary Fig. 5a-b). Interferon gamma response was significantly enriched in non-responders in tumors with immune-low states (NES=-1.30, FDR=0.003, Supplementary Fig. 4a), but not significantly enriched in non-responders in tumors with immune-high states (NES=0.95, FDR=0.33). Interestingly, a set of hypoxia-related signatures, including the Hallmarks hypoxia signature (NES=-1.34, FDR=0.0073), glycolysis (NES=-1.25, FDR=0.0005), epithelial-mesenchymal transition (EMT, NES=-1.21, FDR=0.024), TGF*β* signaling (NES=-1.38, FDR=0.10), angiogenesis (NES=-1.04, FDR=0.23), and VEGF signaling (NES=-1.20, FDR=0.16), were enriched in non-responders with immune-high states (Fig. 3b). In contrast, several of these signatures (hypoxia, glycolysis, and EMT) trended oppositely among non-responders with an immune-low state (Supplementary Fig. 4a).

Further, we calculated a single-sample gene set enrichment score (ssGSEA) for the hypoxia signature in each sample. Uniquely in immune-high samples, ssGSEA scores in samples from patients treated with anti-PD-1 are significantly higher in non-responder compared to responder samples (Fig. 3c, p=0.0051 by Wilcoxon rank-sum). This pattern was not observed in patients with prior anti-CTLA-4 (Supplementary Fig. 4b, n.s. by Wilcoxon rank-sum). This trend was also observed with ssGSEA scores using the glycolysis signature (Supplementary Fig. 4c, p=0.036 by Wilcoxon rank-sum) but not for the epithelial-mesenchymal transition signature or the angiogenesis signature (Supplementary Fig. 4d-e, n.s. by Wilcoxon rank-sum). To affirm that our observations were not overly influenced by the specific set of genes found in the Hallmarks hypoxia signature, we also calculated the ssGSEA score for three other previously published hypoxia signatures [29, 30, 31] and found a significant increase in these signatures in non-responders as well (Supplementary Fig. 4f-h). These data suggest that hypoxia is a biomarker of resistance to anti-PD-1 in metastatic melanoma specifically in immune-high tumors.

### Single cell RNA-seq analysis shows that hypoxia correlates with an immunosuppressive microenvironment in metastatic melanoma

The hypoxic microenvironment contributes to tumor progression and resistance to therapy through multiple mechanisms, including angiogenesis, epithelialmesenchymal transition (EMT), and immune suppression. Hypoxia can impact the TME either directly, through restricting available oxygen [32], or indirectly, via changes in metabolite availability (e.g. lactate, adenosine, etc.), chemical changes (e.g. reactive oxygen species), vascular structure, etc. [33]. To investigate the role of hypoxia in the TME of melanoma, we analyzed a separate cohort of patients with metastatic melanoma with available single cell RNA-sequencing (scRNA-seq). We utilized a subset of pre-treatment samples from the Yang et al. 2024 scRNA-seq cohort (25 samples from 17 patients, Supplementary Fig. 6, additional clinical information in Supplementary Table 7) [24]. We focused our analyses on the pre-treatment scRNA-seq samples to more closely mirror our bulk RNA-seq analyses. After initial quality control filtering, pre-processing, clustering, and removal of doublets and contaminating cells, we had approximately 170,000 cells with cell type annotations at several levels of granularity (Fig. 4a, see details in Supplementary Fig. 7, Methods).

**Figure 4.**
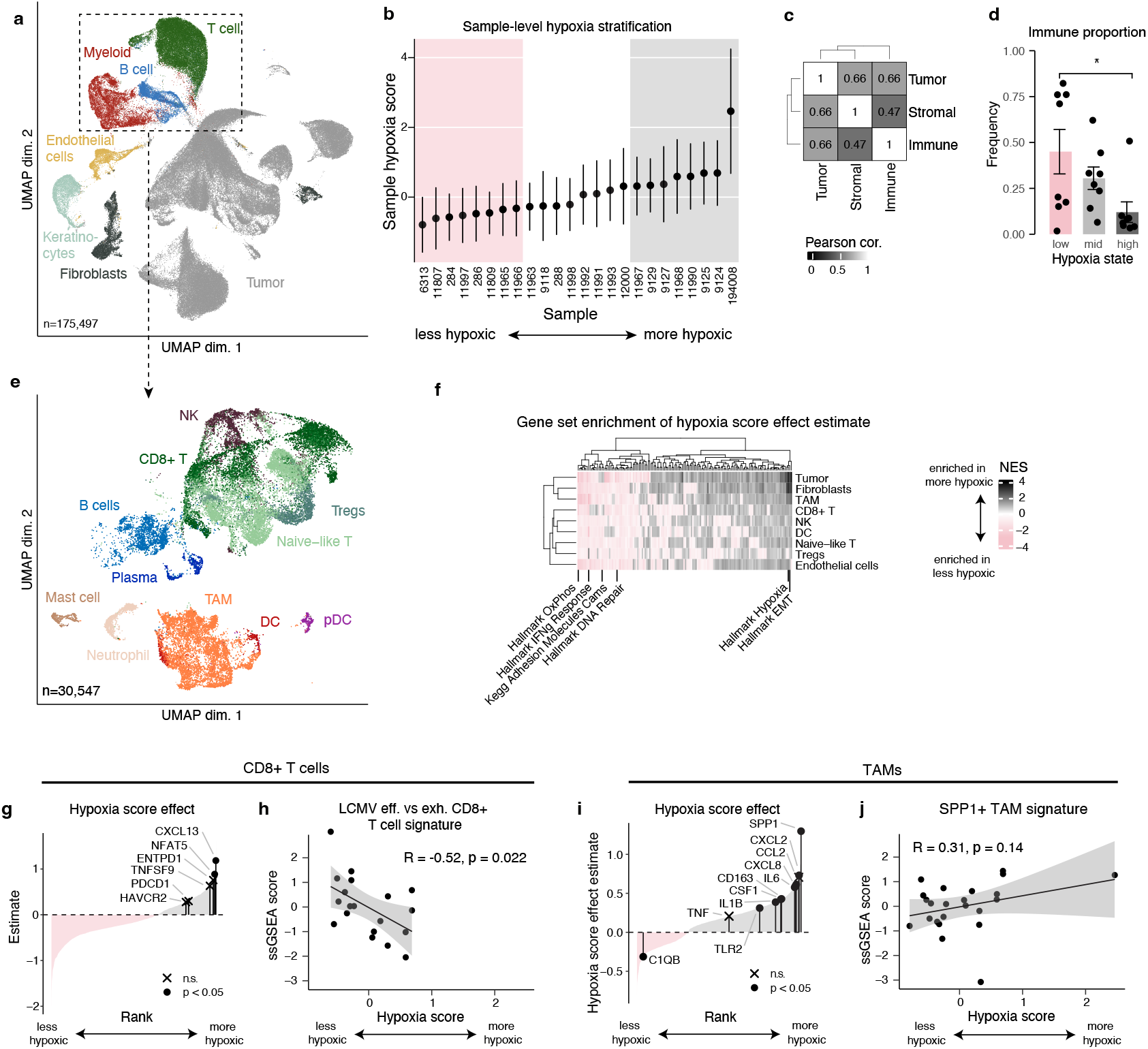
Single-cell RNA-seq reveals hypoxia associates with immune exclusion and immunosuppressive phenotypes. **a**. UMAP showing general cell type clustering for 175,497 cells included in analysis. **b**. Sample-level hypoxia stratification based on a cell-type normalized sample-level hypoxia score (Methods). Samples are designated as low or high based on tertiles. Mean and standard deviation of normalized hypoxia score shown. **c**. Pearson correlation between hypoxia score across compartments. **d**. Immune frequency in each sample by hypoxia state. Significance from Wilcoxon-rank sum, \**p <* 0.05. **e**. UMAP showing immune cell type clustering. **f**. Heatmap showing the pre-ranked GSEA normalized enrichment score (NES) for each of 254 signatures plus 4 additional hypoxia signatures. Genes were ranked by the estimated hypoxia score effect for each cell type as (see Methods on “modeling sample-level hypoxia”). Select gene sets of interest are labelled. Signatures and cell types are clustered by hierarchical clustering. Cell type subsets not shown (B cells, plasma cells, pDCs, mast cells, neutrophils) did not have enough representation across low, mid, and high hypoxia samples to include in the analysis. **g**. Estimated hypoxia score effect for all genes expressed in pseudobulk CD8+ T cells with log2(fold change) greater than 0.1 between high and low hypoxia samples (2,117 genes), as estimated by a linear model. Genes are ranked from lowest to highest based on the estimated effect of the sample-level hypoxia score. Select genes of interest are labelled. **h**. Normalized ssGSEA score of an LCMV effector versus exhausted CD8+ T cell signature in CD8+ T cell pseudobulks correlated with sample-level hypoxia score (Pearson’s *R* = −0.52, *p <* 0.05). Note that sample 194008 (hypoxia score of 2.46), is dropped from this panel because no CD8+ T cells were identified from this sample. **i**. Same as **g**. for tumor-associated macrophages (TAMs, 2,107 genes). **j**. Normalized ssGSEA score of the SPP1+ TAM signatures from Wei et al. 2021 [43] in TAMs correlated with sample-level hypoxia score (Pearson’s *R* = 0.31, *p* = 0.14).

We then calculated a “hypoxia score” for each sample (Methods). After scoring each cell for the Hallmarks hypoxia signature (per-cell values can be found in Supplementary Table 8), we created high, mid, and low hypoxia categories by dividing the cohort by tercile of the mean hypoxia score per sample (Fig. 4b). Then, to evaluate whether the levels of the hypoxia signature were driven by a global cellular response to an environmental hypoxia or a melanoma-intrinsic pseudohypoxia [34, 35], we examined the correlation of the mean levels of hypoxia signature expression between the tumor, immune, and stromal compartments. The hypoxia signature was robustly correlated between the tumor cells and other compartments (Pearson’s correlation of 0.66, Fig. 4c), which is more consistent with a global cellular response to hypoxia. More hypoxic samples also tended to be immune excluded (Fig. 4d), consistent with our expectations [36].

To investigate changes in the immune milieu, we subset the immune cells, defined by high expression of *PTPRC* (CD45), and performed dimensionality reduction, clustering, and cluster annotation to identify immune cell subsets (Fig. 4e), Supplementary Fig. 7b). Subtle shifts in the composition of the immune microenvironment, both in more hypoxic and immune excluded samples were apparent but not significant in this cohort (Supplementary Fig. 8a-f). Then, we tested the relationship between immune cell type genes most associated with overall sample levels of hypoxia using a pseudobulk approach. We first fit linear models predicting gene expression from the hypoxia score for each gene (Supplementary Table 9). We then performed a similar pre-ranked gene set enrichment analysis as with the bulk RNA-seq analysis, utilizing the same 254 gene sets with four hypoxia-related signatures. Hierarchical clustering of the signature scores revealed a cohesive set of signatures which broadly associated with sample-level hypoxia (Fig. 4f). Across all cell types in more hypoxic samples, we observed upregulation of hypoxia response and EMT genes, and downregulation of oxidative phosphorylation (consistent with a progressive loss of mitochondria and oxidative metabolism in a hypoxic TME [37]), interferon gamma response, adhesion, and DNA repair.

Within CD8+ T cells, we found that higher sample-level hypoxia associated with higher levels of *CXCL13* (a marker which has been shown to have discriminating power for antigen specificity in CD8+ T cells in the TME [38]) and *ENTPD1* (CD39), which have been previously associated with CD8+ T cell exhaustion in the TME [39, 40] (Fig. 4g). Accordingly, an LCMV signature for effector versus exhausted T cells also decreases as hypoxia increases (Fig. 4h, additional exhaustion signatures shown at Supplementary Fig. 8g-h) [41, 42]. We also found that the tumor-associated macrophage (TAM) cluster had high levels of immuno-suppressive markers, such as *CXCL8, CSF1, CD163*, and *SPP1* (Fig. 4i-j, additional TAM signatures shown at Supplementary Fig. 8i-j), which has been found to be upregulated in tumor-associated macrophages in previous studies [43]. More pro-inflammatory TAM markers, such as *C1QB* and *TNF* were not as strongly associated with hypoxia. Overall, we observed a depletion of immune cells in more hypoxic samples, as well as a potentially more immunosuppressive microenvironment.

Further, we found similar trends in two independent, previously published scRNA-seq cohorts of patients with metastatic melanoma, Jerby-Arnon et al. 2018 (Supplementary Fig. 9a-d) [44] and Sade-Feldman et al. 2018 (Supplementary Fig. 10a) [11]. Since the Sade-Feldman dataset did not contain tumor or stromal cells, the analysis was reduced and adapted (see Methods for details). Looking only at untreated metastatic samples, we performed a similar hypoxia stratification (Supplementary Fig. 9e-g and 10b) and found that hypoxia also associated with immune exclusion (Supplementary Fig. 9h). Despite the lower cell counts in these datasets, we observed similar expression patterns of *EPAS1, FLT1, HIF1A, KDR*, and *VEGFA* across tumor, stromal (fibroblasts, endothelial cells), and immune cells (T cells, myeloid cells) in these cohorts (Supplementary Fig. 9i and 10e). Although the enrichment of the specific gene signatures varied in this external dataset, we observed a similar enrichment of a more terminally exhausted phenotype in CD8+ T cells and a more immunosuppressive phenotype in the TAM population (Supplementary Fig. 9j-o and 10f-h).

### Hypoxia associates with unique stromal-tumor interactions

We hypothesized that the impact of hypoxia on non-immune cells, like cancer-associated fibroblasts and endothelial cells (ECs), could also play a role in resistance to checkpoint blockade. The hypoxia score was highest in tumor and stromal cells (keratinocytes, ECs, and fibroblasts, *p <* 0.05 for each group in one versus rest Wilcoxon rank-sum, Fig. 5a). The expression of the key hypoxia-sensing transcription factors *HIF1A* and *HIF2A* as well as the downstream angiogenesis-related genes *VEGFA, FLT1* (VEGFR1), and *KDR* (VEGFR2) were also high in these subsets (Fig. 5b). *HIF1A* was expressed across all cell types, but was significantly higher in ECs, fibroblasts, and myeloid cells. In contrast, *HIF2A* (*EPAS1* gene) was primarily expressed in ECs and fibroblasts. *VEGFA* was significantly high in tumor cells, fibroblasts, and myeloid cells. Fibroblast frequency was associated with higher hypoxia (Fig. 5c), and they also showed increased expression of certain chemokines and cytokines in more hypoxic samples (Fig. 5d). Sample-level hypoxia did not correlate with EC frequency (Fig. 5e). However, in more hypoxic samples, ECs expressed genes promoting angiogenesis, whereas in less hypoxia samples, ECs expressed genes promoting adhesion and certain chemokines, like *CXCL10* (Fig. 5f).

**Figure 5.**
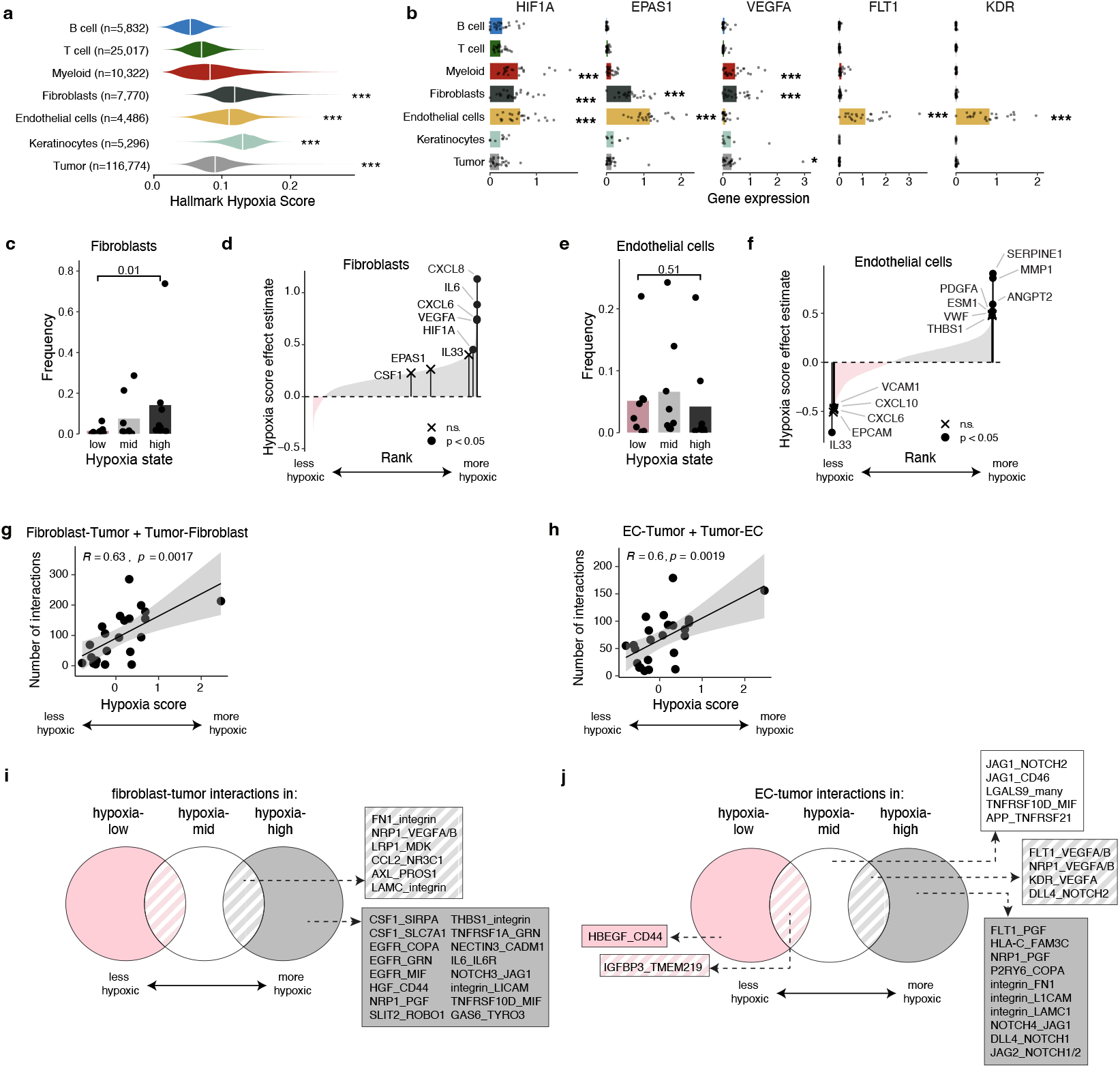
Effect of hypoxia on stromal cell characteristics and stromal-tumor communication. **a**. AUCell Hallmark Hypoxia score for all cells in each general cell type. **b**. Log-normalized expression (counts per ten thousand) of labelled gene for each cell type averaged across all cells of that type per sample. **a-b**. Cell types with significantly higher scores by Wilcoxon rank-sum are shown, with each cell type compared against all others. *: *p <* 0.05, **: *p <* 0.01, ***: *p <* 0.001. **c**. Frequency of fibroblasts (out of all cells) in low-, mid-, and high-hypoxia samples. **d**. Estimated hypoxia score effect for all genes expressed in pseudobulk of labelled cell type with log2(fold change) greater than 0.25 between high and low hypoxia samples (8,410 genes in fibroblasts), as estimated by a linear model (see Methods for details). Genes are ranked from lowest to highest based on the estimated effect of the sample-level hypoxia score. Select genes of interest are labelled. **e**. Frequency of endothelial cells (out of all cells) in low-, mid-, and high-hypoxia samples (5,871 genes in endothelial cells with log2(fold change) greater than 0.25. **f**. Like **d**., but with endothelial cells. **g-h**. The relationship between the number of stromal-tumor interactions estimated by the CellPhoneDB statistical analysis method [45] and the sample-level hypoxia score for fibroblasts and endothelial cells (ECs). Correlations shown are Pearson correlations. **i-j**. All stromal-tumor interactions which are significant ≥ 75% of the samples (*p <* 0.05 in at least 6/8) in each category: low hypoxia (left, pink), mid hypoxia (middle, white), high hypoxia (right, grey). All ≥ 75% significant interactions in multiple categories (e.g. ‘KDR_VEGFA’ in EC-tumor interactions for both mid hypoxia and high hypoxia) are shown in dashed overlapping regions. Interacting pairs are listed as “partnerA_partnerB”, where partnerA is expressed on stromal cells and partnerB is expressed on tumor cells.

Given the high expression levels of hypoxia- and angiogenesis-related transcription factors, as well as the phenotypic changes in stromal cells, we wanted to understand if and how stromal cells could be interacting with tumor cells. So, we inferred cell-cell interactions based on the gene expression of receptor-ligand pairs using CellPhoneDB [45] (Supplementary Table 10). The number of significant stromal-tumor interactions positively correlated with hypoxia score for both ECs and fibroblasts (Fig. 5g-h). Other interactions, such as the frequency of stromal-immune interactions, were largely not correlated with the sample-level hypoxia score (Supplementary Fig. 11a-b). We then examined stromal-tumor interactions consistently found only in hypoxic samples by identifying interactions found in *≥* 75% of the samples (i.e. 6/8) in a particular hypoxia tertile (high, mid, or low; Fig. 5i-j). Fibroblasts in the high- and mid-hypoxia groups were identified to consistently have FN1 (fibronectin)-integrin and NRP1-VEGF interactions with tumor cells. However, several fibroblast-tumor interactions were only observed in the high-hypoxia group: fibroblast expression of EGFR interaction with COPA, GRN, and MIF on tumors, as well as CSF1 expression on fibroblasts interacting with markers expressed on tumors. ECs in mid- and high-hypoxia samples also had NRP1-VEGF interactions, in addition to many angiogenic signals downstream of HIF1A and HIF2A, including FLT (VEGFR1) to VEGFA/B, KDR (VEGFR2) to VEGFA. Unique to the ECs in high-hypoxia samples were interactions involving ECs expression of integrins as well as an increase in EC expressed DLL4 and JAG2 to tumor expressed NOTCH receptors. Altogether, these signals reveal a concert of stromal interactions which may be reinforcing or responding to hypoxia in the melanoma TME and subsequently promoting angiogenic and immunosuppressive signals.

### Targeting HIF-2*α* signaling in combination with anti-PD-1 in pre-clinical models improves tumor control

We hypothesized that more proximal inhibition of hypoxia-associated pathways may lead to greater clinical benefit. Based on our single-cell RNA-seq analysis, we identified *EPAS1* (HIF-2*α*) as being highly and selectively expressed on endothelial cells and fibroblasts within the TME (Fig. 5b, Supplementary Fig. 9i). HIF-2*α* inhibitors have recently been FDA-approved for clear cell renal cell carcinoma (ccRCC), a tumor which expresses HIF-2*α*. On the other hand, melanoma, along with infiltrating immune cells, expresses HIF-1*α* (Fig. 5b). In the treatment of melanoma, we therefore reasoned that inhibiting HIF-2*α* may operate indirectly, specifically targeting stromal and endothelial cells (as opposed to tumor cells) without impairing immune effector function. Moreover, we hypothesized that inhibiting HIF-2*α* would be more effective in immune-high tumors.

To test this hypothesis, we used a syngeneic murine melanoma tumor model, B16.F10 (B16). The tumor microenvironment of B16 recapitulates several features that we observed from the human clinical setting in our scRNA-seq analysis. We analyzed a previously published dataset of bulk RNA-sequencing of B16 tumors (GSE96972) [46] and found that, similarly to the human clinical context, B16 tumors expressed very low levels of *Epas1* (HIF-2*α*, Supplementary Fig. 12a). Additionally, from previously published single-cell sequencing of the TME of B16 tumors, *Hif1a* was broadly expressed across immune and stromal cells, but only fibroblast and endothelial cells had relatively higher expression levels of *Epas1* (Supplementary Fig. 12b). Then, to compare the immune-high and immune-low settings, we also utilized B16.F10 expressing the model antigen ovalbumin (B16-OVA). Respectively, B16 and B16-OVA represent a poorly immune-infiltrated tumor that is unresponsive to anti-PD-1 alone and a more highly immune-infiltrated tumor that is partially responsive to anti-PD-1 [47, 48, 49]. Mice were implanted with B16 or B16-OVA tumors, and treated with either anti-PD-1, a small molecule inhibitor of HIF-2*α*, PT2399, or the combination of anti-PD-1 with PT2399 (Fig. 6a). Mice implanted with B16-OVA tumors and treated with the combination anti-PD-1/PT2399 showed delayed tumor growth relative to control or each treatment administered alone (Fig. 6b). In contrast, mice implanted with the relatively immune-excluded B16 parental tumor did not see significantly improved tumor growth control when treated with the combination anti-PD-1/PT2399 (Fig. 6c). This suggests that the benefit of HIF-2*α* inhibition occurred in combination with immunotherapy and in tumors more sensitive to anti-PD-1.

**Figure 6.**
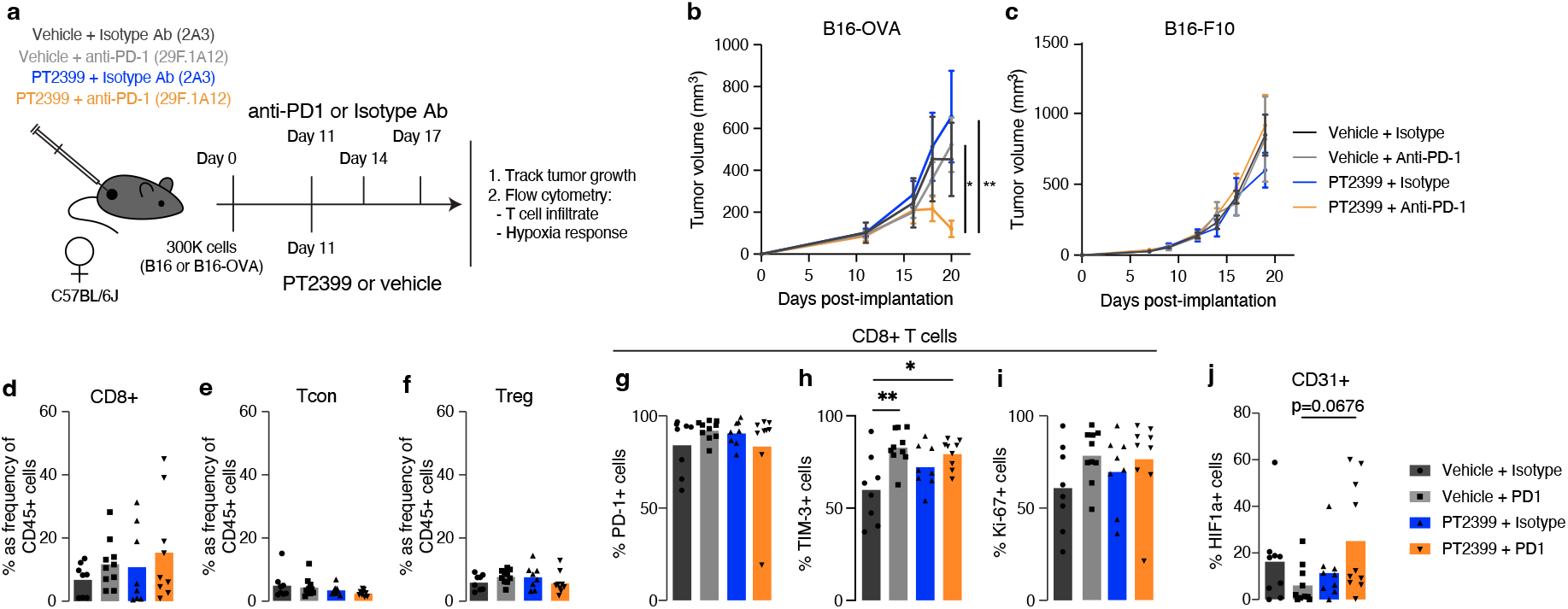
Targeting hypoxia in combination with anti-PD-1 slows tumor growth in pre-clinical models of immune-high tumors. **a**. Graphic depicting experimental design of combination PT2399/anti-PD-1 treatment experiments. **b-c**. Growth curves showing tumor growth in response to HIF2*α* inhibitor (PT2399) and anti-PD-1 blockade combination treatment for (b) B16-OVA and (c) B16.F10. Data representative of two independent experiments (*n >* 5 per group). **d-i**. Flow cytometry characterization of T cells isolated from B16-OVA tumors treated with isotype control, anti-PD-1, PT2399, or the combination. Data showing two independent experiments combined (n=4-5 per group). **d**. Percentage of CD8+ T cells of CD45+ cells. **e**. Percentage of conventional T cells (Tcon; CD4+FOXP3-) of CD45+ cells. **f**. Percentage of regulatory T cells (Tregs; CD4+FOXP3+) of CD45+ cells. **g**. Percent of PD-1+ CD8+ T cells out of total CD8+ T cells. **h**. Percent of TIM-3+ CD8+ T cells out of total CD8+ T cells. **i**. Percent of Ki-67+ CD8+ T cells out of total CD8+ T cells. **j**. Percentage of HIF-1*α*+ CD31+ cells out of total CD31+ cells in the tumor. Significance levels are denoted as \**p <* 0.05, \*\**p <* 0.01, \*\*\**p <* 0.001.

To understand the immunologic mechanism by which the combination anti-PD-1/PT2399 exerted its effects, we conducted cellular analyses of T cells isolated from B16-OVA tumors on day 4-5 after initiation of treatment (Supplementary Fig. 13, 14, 15). We observed no significant numerical advantage in CD8+ T cells or decrease in regulatory T cells (Fig. 6d-f). Furthermore, despite differences in tumor growth, we saw no difference in CD8+ T cell phenotype or function, including co-inhibitory receptors (PD-1, TIM-3), proliferation (Ki-67), or granzyme B expression between the anti-PD-1 treated group and the anti-PD-1/PT2399 group (Fig. 6g-i, Supplementary Fig. 12c). Likewise, we saw no differences in the CD8/Treg ratio (Supplementary Fig. 12d) or the regulatory phenotype based on PD-1, Ki-67, or CD25 expression (Supplementary Fig. 12e-g). Given the high expression of HIF-2*α* in fibroblasts and endothelial cells relative to melanoma or immune cells (Fig. 5b, Supplementary Fig. 9i), we suspected that the HIF-2*α* function may be greater on stromal and endothelial cells, and that re-modeling of the tumor microenvironment could be responsible for slowing tumor growth, and also performed flow cytometry on CD45-CD31+ cells, which represent endothelial cells and some stromal cells (Supplementary Fig. 16 and 17). Consistent with this hypothesis, intra-tumoral CD31+ cells showed an increase in HIF-1*α* expression in anti-PD-1/PT2399 treated mice, demonstrating that non-immune cells are impacted by the combination therapy (Fig. 6j). These findings suggest that combined anti-PD-1/HIF-2*α* inhibition may compensate for the lack of HIF-2*α* signaling by up-regulating HIF-1*α* on non-immune cells in the tumor microenvironment, and thereby improve tumor growth control.

## Discussion

In this study, we performed a meta-analysis of all previously published cohorts of patients with advanced melanoma treated with anti-CTLA-4, anti-PD-1, and combination anti-CTLA-4/anti-PD-1 for whom clinical annotations and bulk RNA-seq data were available. In stratifying patient populations by immune state, we identified several important patterns of response between treatment groups. First, we recapitulated the observation that an immune-high microenvironment predicted response to anti-PD-1 only in prior anti-CTLA-4 treated patients [18], highlighting the confounding effects of prior immunotherapy on biomarkers of response. We emphasize that features of response and non-response are sufficiently distinct for patients with and without prior anti-CTLA-4 that they ought to be considered separate populations; to this end, we have provided an updated, organized data table for all cohorts included in our analysis as a resource for future analyses (Supplementary Table 1). Second, we found that the increased response rates of combination blockade over single agent anti-PD-1 occurs in tumors classified as immune-high, but not immune-low, supporting a “rich-get-richer” scheme for response to combination blockade. Our analysis suggests that patients with immune-high tumors may be the specific incremental beneficiaries of combination blockade, and that, counter-intuitively, toxicity could be decreased while maintaining efficacy by giving anti-PD-1 monotherapy to patients with immune-low tumors. Additional work would certainly be required to translate our transcriptomic metric of immune infiltrate into the clinic, though the rapid development and increasing power of computer vision models to quantify and classify features from hematoxylin and eosin (H&E) staining [50] or other common clinical assays provides a potential path forward.

Our immune-high versus immune-low classification also provides a framework in which to separate mechanisms of resistance to ICB. An immune-high microenvironment suggests that the immune response has been partially successful: immune cells have infiltrated, but the anti-tumor killing process has been stymied. While previous clinical analysis have suggested hypoxia broadly associates with resistance to ICB [9] and other solid tumors [51], our analysis demonstrated an associated of hypoxia with poor response that is specifically enriched in immune-high tumors and underscores the importance of heterogeneity within the TME. Our data therefore suggest that therapies targeting hypoxia in the tumor microenvironment may be more effective in highly immune-infiltrated tumors.

In melanoma, clinical trials targeting angiogenesis alone have been met with relatively little success [52]. In more recent years, there have been several trials combining hypoxia inhibitors or anti-angiogenic therapeutics with immune-checkpoint blockade [53, 54], demonstrating some promise for the combination [55, 56]. However, the LEAP-003 trial [57], testing lenvatinib (multiple tyrosine kinase inhibitor whose targets include VEGF receptors) and pembrolizumab (anti-PD-1) as a first-line combination therapeutic and focusing on overall survival (OS) outcomes, was discontinued due to a lack of improvement in OS relative to anti-PD-1 alone [58]. Given our results, it is possible that an immune-stratified analysis of previous, ongoing, and upcoming trials involving combination angiogenesis inhibitors and ICB could enrich for a responding patient population.

While hypoxia is considered a general feature of solid tumors, in our scRNA-seq analysis of pre-treatment metastatic melanomas, we observed varying levels of hypoxia response in different samples (Fig. 4b, Supplementary Fig. 9f and 10). Differences in levels of hypoxia then corresponded to varying degrees of the immunosuppressive and tissue remodeling impacts of hypoxia. Hypoxia was associated with decreased immune infiltration overall, and the immune cells which were present had different phenotypes: T cells tended to be more terminally exhausted [42] and had higher levels of *ENTPD1* (CD39), a transcription factor which promotes suppressive activity in exhausted CD8+ T cells [41]. Macrophages possessed an SPP1+ phenotype previously associated with immunosuppression [43]. In addition to immune cells, we observed that stromal cells had the highest hypoxia response scores, suggesting an important role of stromal cells in mediating hypoxia effects in the microenvironment. In more hypoxic samples, fibroblasts and endothelial cells expressed genes which could contribute to remodeling the TME (e.g. *CXCL8, IL6*) and had increased numbers of stromal-tumor interactions. Overall, our analysis paints a picture of high variance in hypoxia levels in melanomas and a subsequent differential impact on immunosuppression and tumor-stromal signaling, potentially promoting tumor growth and favoring an immunosuppressive microenvironment.

Within our scRNA-seq data, we also identified differential gene expression of commonly clinically targeted gene products across cell types. For example, *VEGFA* (VEGF; targeted by lenvatinib) was expressed on myeloid cells and fibroblasts and to a lesser extent on keratinocytes and tumor cells; *KDR* (VEGFR2; targeted by bevacizumab, a VEGFR2 blocking antibody) on endothelial cells; and *EPAS1* (HIF-2*α*; targeted by belzutifan, the human analog to PT2399) expressed on endothelial cells and fibroblasts (Fig. 5). The differential expression of HIF-2*α* in fibroblasts in addition to endothelial cells may also offer a unique and novel target.

Fibroblasts are enriched in more hypoxic melanomas (Fig. 5i, Supplementary Fig. 8h) and can reinforce hypoxia in the microenvironment through collagen deposition and increased fibrosis [59]. The HIF-2*α* inhibitor belzutifan is already approved to treat von Hippel-Lindau (VHL) disease-associated tumors [60] and for clear cell renal cell carcinoma (ccRCC) in combination with a PD-1 or PD-L1 inhibitor and a vascular endothelial growth factor tyrosine kinase inhibitor (VEGF-TKI) [61]. In these tumor types, belzutifan is thought to act on a tumor-intrinsic features, since these tumor types are known to commonly mutate HIF-2*α* [62]. HIF-2*α* is not commonly expressed by melanomas, but the high expression levels seen in stroma offer a potential opportunity to normalize tumor vasculature, which is a previously proposed mechanism of anti-angiogenic therapies [63]. Additionally, although HIF-2*α* expression has also been preferentially shown to be expressed in regulatory T cells, as opposed to HIF-1*α* in CD8+ T cells [64, 65, 66], alterations in regulatory and CD8+ T cell phenotypes were not seen in our pre-clinical models, suggesting that the impact of HIF-2*α* inhibition is more indirect on T cell functionality. In all, our data is consistent with the addition of HIF-2*α* inhibition acting through an indirect, stromal-focused mechanism, thus marking a novel approach. Overall, our study emphasizes the utility of curating multiple, large, high quality, clinically annotated datasets into a larger cohort. Clinical datasets oftentimes contain patients with varied treatment histories and heterogeneous biology, making the interpretation of these datasets very difficult. However, given this heterogeneity, collecting larger cohorts with organized annotations will be crucial to achieving the number of samples required to derive meaningful biological insights. We demonstrate how careful stratification by clinical and molecular characteristics can be leveraged to derive meaningful biological insights and lead to the rational discovery of novel clinical targets for combination therapy.

## Declarations

## Acknowledgements

We thank past and present members of the Sharpe, Van Allen, and Liu laboratories for thoughtful scientific discussions. We are grateful to Dirk Schadendorf and Bastian Schilling for clarifications about clinical annotations in the Liu 2019 cohort, James Wilmott and Tuba Gide for their correspondence and clarification of clinical features of the 2019 Gide cohort, and Katie Campbell and Antoni Ribas for their correspondence and clarifications of clinical features of the CheckMate067 subset of the 2023 Campbell cohort. This work was supported by the American-Italian Cancer Foundation Post-Doctoral Research Fellowship (Giuseppe Tarantino), the Society for Immunotherapy of Cancers (David Liu), the Doris Duke Charitable Foundation Clinical Scientist Training Program (David Liu), the National Institutes of Health (NIH K08CA234458, David Liu), the Eleanor and Miles Shore Faculty Development Award (David Liu), the Renal SPORE (NIH P50CA101942, Arlene Sharpe, Kelly Burke), the Barr Foundation (Kelly Burke), the Adelson Medical Research Foundation (Genevieve Boland), Bill and Emma Roberts MGH Research Scholar Program (Genevieve Boland), and the Patricia K. Donahoe Award from the Huiying Foundation (Genevieve Boland).

## Ethics approval

This retrospective study and associated informed consent procedures were approved by the central Ethics Committee (EC) of the Dana Farber Cancer Institute (IRB 05-42). Approval by the local EC was obtained by investigators if required by local regulations. All pre-clinical experiments using mouse models were performed at Harvard Medical School in accordance with ethical regulations approved by the Harvard Medical Area Standing Committee on Animals and the American Association of Accreditation of Laboratory Animal Care.

## Code availability

Python and R code is available in packages as described in the manuscript. Code to regenerate all analyses and figures in a Quarto book is available at GitHub at https://github.com/davidliu-lab/diffIO-hypoxia-figures. Additional reasonable requests for code will be promptly reviewed by the senior authors to verify whether the request is subject to any intellectual property or confidentiality obligations and shared to the extent permissible by these obligations.

## Data availability

Raw data from all previously published work can be obtained by contacting corresponding authors of cited works. All analyzed or processed data included in the bulk meta-analysis and the single-cell analyses are in supplementary tables or as .rds objects available on GitHub at https://github.com/davidliu-lab/diffIO-hypoxia-figures. Raw FASTQ files for additional bulk RNA-seq samples for the 2024 Yang cohort are available on GEO (accession number GSE274749).

## Authors’ contributions

A.Y.H., K.P.B., A.H.S., and D.L. conceived and designed the overall study. K.P.B., R.P., L.M., P.F., and V.R. performed experiments and data analysis. A.Y.H., N.V., C.R., E.R., G.T., T.J.A., M.G., J.C., Y.H., J.Y., D.F., and C.W. acquired and processed data, and/or performed select data analysis. L.L.H. and K.G. performed sample processing; G.M.B. and M.K. oversaw sample processing. G.F. contributed key resources. A.Y.H., K.P.B., E.I.B., F.S.H., G.M.B., M.K., A.H.S., and D.L. interpreted the data. A.Y.H., K.P.B., and D.L. wrote the manuscript. All authors reviewed and edited the manuscript.

## Methods

### Patient cohorts for bulk RNA-seq analysis

For the analysis, pre-treatment samples of tumors from patients with metastatic melanoma treated with immune checkpoint blockade were identified from previously published work. Samples from cohorts with fewer than 10 samples total were not considered (e.g. Hugo 2016 [9]). The following cohorts were included in the meta-cohort: Van Allen 2015 (n=156) [6], Weber 2016 (n=59) [21], Riaz 2017 (n=80) [22], Liu 2019 (n=169) [18], Gide 2019 (n=74) [16], Freeman 2022 (n=59) [23], Campbell 2023 (n=59) [19], which contains patient data from patients described in [4], and Yang 2024 (n=37) [24]. Clinicopathological and demographic data were initially obtained from the published papers. Additionally, we contacted authors of the previously published work for additional information on prior therapies and updated survival data (Supplementary Table 1).

### Clinical exclusion criteria

Only cutaneous melanoma was considered in our analysis. Tumor types of unknown origin were excluded. Additionally, some samples had complex treatments given concurrently with ICB (e.g. tyrosine kinase inhibitors) and those patients were excluded.

### Definitions of response

Response was evaluated either by RECIST v1.1 where available, or by best overall response (BOR). Responders (R) were categorized by having RECIST or BOR complete or partial response (CR, PR) or stable disease (SD) with PFS greater than six months. Non-responders (NR) were categorized by having RECIST or BOR progressive disease (PD) or stable disease (SD) with PFS less than six months. Since the interpretation is less clear, patients with mixed response (MR) or non-evaluable response (NED) were excluded in all analyses involving response rates. For the Weber 2016 cohort, samples were taken from patients who were treated sequentially with anti-PD-1 and anti-CTLA-4; for these samples, we only considered response to the first line of treatment as response to the monotherapy.

### Inclusion in survival analysis

Because patients in the Weber 2016 cohort were treated sequentially with anti-PD-1 and anti-CTLA-4, samples from the Weber 2016 cohort were excluded from all survival analysis (n=59). Since several samples from the Freeman 2022 cohort also had subsequent treatment, we also removed them from the survival analysis (n=59). All samples from the 2023 Campbell (067) cohort were missing censoring data, and thus they were also excluded from survival analysis (n=59).

### Bulk RNA-sequencing processing and analysis of clinical cohorts

#### Alignment and quantification

To harmonize the data, we obtained raw FASTQ files for all previously published bulk RNA-sequencing samples. We evaluated each FASTQ files’ quality through examining metrics from FASTQC (v0.11.9). We utilized STAR (v2.7.0) [67] to align samples to the hg19 reference genome, producing BAM files. BAM files were quantified using salmon (v0.14.1) [68]. All alignment and quality control methods were run utilizing the Terra platform for biomedical research.

#### Quality control filtering

For quality control, we evaluated STAR alignment metrics, removing samples with fewer than 1 million reads and/or fewer than 50% uniquely mapped reads. Only protein coding genes expressed in at least 5% of all samples were included. We then regenerated a new TPM metric for each sample to normalize the total transcriptome sum to 1 million. Additional outliers were identified through principal component analysis (PCA) of the STAR metrics and gene expression pre- and post-batch correction.

#### Batch correction

We observed a batch effect capture in the first principal component of the non-batch corrected data that was largely explained by sequencing modality (i.e. poly-A tail capture versus transcriptome capture technology; Supplementary Fig. 2b-c). We utilized ComBat-Seq [69], implemented in the R package sva (v3.46.0), to correct for the batch effect by sequencing modality. We chose not to batch correct by cohort because we reasoned that there could be relevant biological differences between cohorts, which ought to be preserved (Supplementary Fig. 2d-e). We again regenerated a new TPM metric for each sample to normalize the total transcriptome sum to 1 million.

#### Immune stratification and calculation of the immune score

To characterize the TME, we utilized sample-level signature scoring to categorize samples. Each sample was scored using ssGSEA for cell type/microenvironmental signatures, defined in Bagaev et al. 2021 [25]. We observed that several of the signatures were highly co-correlated; hierarchical clustering of the co-correlation matrix divided the signatures into three sets: a set of 19 immune signatures, 8 stromal/granulocytic signatures, and 2 outliers related to trafficking and proliferation (Supplementary Fig. 3a).

We calculated the immune score from the 19 immune signatures. For each signature, we scaled and centered such that the values were z-scored. Then, the mean across all signatures for each sample is the immune score, which was subsequently also z-scored.

Given high levels of correlation between immune signatures, we chose to categorize samples by their overall level of immune signatures, as opposed to the signature scores of specific cell types. Thus, we hierarchically clustered samples by their levels of the immune signatures. Clusters were assigned based on overall expression levels of the immune signatures, with the samples in the cluster with higher expression of immune-related signatures defined as “immune-high” and the samples in the cluster with lower expression of immune-related signatures defined as “immune-low”. For each sample, we also calculated an “immune score” which is defined as the mean of the scaled ssGSEA scores for each of the immune signatures.

To understand how this signature related to preexisting signatures which predict response to ICB, we also calculated scores for nine previously published immune-related signatures: (1) expanded IFN*γ* signature [12], (2) Hallmark IFN*γ* signature [26], (3) TLS marker genes [27], (4) CIBERSORTx Tirosh immune proportion [70], (5) CIBERSORTx LM22 absolute immune score [71], (6) non-expanded IFN*γ* signature [12], CIBERSORTx LM22 CD8+ T cell proportion [71], CIBERSORTx Tirosh CD8+ T cell proportion [70], (9) Cabrita TLS signature [27]. For (1), (2), (3), (6), (9), we calculated scaled ssGSEA scores from our samples. For (4), (5), (7), (8), we used the referenced signature matrix and CIBERSORTx to deconvolute the cell type proportions from the bulk RNA-seq samples.

Association between discrete clinical variables and the immune score were calculated using Fisher’s Exact Test. For biopsy site versus immune state, the number of immune-high and immune-low tumors from lymph node (n=34 total; 12 immune-high, 6 immune-low, and 16 without RNA-seq), skin (n=141; 68 immune-high, 36 immune-low, and 37 without RNA-seq), and other biopsy sites (n=41; 12 immune-high, 9 immune-low, and 20 without RNA-seq) were considered for the contingency matrix. Any samples with unknown biopsy site annotations were not included in the contingency matrix. For antigen presentation mutations versus immune state, the number of immune-high and immune-low tumors with (38 immune-high, 25 immune-low) or without (67 immune-high, 40 immune-low) antigen presentation mutations were considered for the contingency matrix. The following genes were considered as antigen presentation genes: *RFXANK, RFXAP, RFX, CIITA, NLRC5, B2M, HLA-C, HLA-C, HLA-A, ICAM1, PTK2, SCIN, EFNA1, TAP1, TAP2, TAPBP, PDIA3, CALR, CANX, HSPA5, TPP2, ERAP1, ERAP2, PSMB10, PSMB9, PSMB8*. Silent, intron, 3’UTR, and 5’UTR mutations were not considered in the count.

#### Multi-variable model

We constructed a multi-variable logistic regression model to predict response from treatment and the immune score or the sample immune state (model formulation: response ∼ immune_score + treatment). For the treatment variable, we used the anti-PD-1 treatment group as the reference. To affirm that our model was appropriate and parsimonious, we also tested models with fewer variables (e.g. response ∼ treatment, response ∼ 1) and compared the Akaike information criteria (AIC) between the models.

#### Differential gene expression (DGE) analysis

DGE analysis was performed to identify genes differentially expressed between responders and non-responders for each treatment group and immune state. For the DGE analysis, to best capture features which are characteristic of response or non-response to ICB treatment, we did not include patients with response annotations of SD. For each gene, we calculated log2 fold change of transcripts per million (TPM) between responder samples and non-responder samples. Given that Wilcoxon rank-sum is best at FDR control when analyzing human population samples [72], we calculated the significance of the difference in TPMs by Wilcoxon rank-sum, utilizing functions from the tidyverse (v2.0.0) collection of packages in R.

#### Pre-ranked gene set enrichment analysis

Gene signatures were obtained using the msigdbr (v7.5.1) package in R, which accesses the Molecular Signatures Database [73]. 254 gene signatures were selected for the analysis, including: 50 signatures from the Hallmarks database [26], 186 signatures from the Kegg database [74], and 18 selected signatures from ImmuneSigDB [75]. For additional analyses, we also examined 4 hypoxia-related signatures from C2: ‘BUFFA_-HYPOXIA_METAGENE’, ‘HARRIS_HYPOXIA’, ‘KIM_-HYPOXIA’, and ‘WINTER_HYPOXIA_METAGENE’.

To understand gene signatures enriched in responders and non-responders for each treatment group and immune state, pre-ranked GSEA was calculated using the Lightweight Iterative Gene set Enrichment in R (LIGER) tool (v2.0.1). Genes were ranked by the signed -log10(p-value) estimated from the DGE analysis comparing responders and non-responders, where the direction of the value was determined by the log2 fold change.

For per-sample gene signature scoring, single sample GSEA (ssGSEA) signature scoring [76] was performed on the TPM expression of all genes in a sample using the GSVA package (v1.46.0).

To identify gene signatures which were differential between immune-high and immune-low for each treatment group, we calculated a “differential enrichment score”, which was the difference between the normalized enrichment score (NES) of each signature in immune-high versus immune-low. Then, we ranked all the gene signatures by their score from highest to lowest.

#### Genomic analysis

For samples with available paired whole-exome sequencing, we re-analyzed samples with the Broad Institute CGA pipeline, using the TERRA platform, adopting the same quality control filters used for Liu et al. Nature Medicine 2019 [18]. We estimated tumor purity, ploidy, and individual mutations through FACETS [77]. We calculated the number of clonal, sub-clonal, and the number of non-synonymous mutations. Tumor mutational burden was defined as the number of non-synonymous mutations divided by the mean tumor target coverage.

### Single cell RNA-seq dataset processing and analysis of Yang et al. 2024

This section will describe the sample processing and analysis for subset of samples from the Yang et al. 2024 dataset [24] which were utilized in this study.

#### Sample selection

The sample collection and processing of the single cell cohort is described in Yang et al. 2024 [24]. Initial data processing of the complete dataset was performed using the scanpy software in Python [78]. From the dataset, we selected a subset of 24 pre-treatment samples from 17 patients. Within this subset, there were 177,737 cells total. Of these, there were 116,774 tumor cells, 43,411 immune cells, and 17,552 stromal cells (“compartments” as shown in Supplementary Fig. 7a). Details of their cell type identification can be found in Yang 2024. Data for all cells in each compartment (tumor, immune, stromal) were then re-processed using the Seurat package (v3.1.1) in R (v4.2.1) [79]. Gene expression measurements were normalized and transformed using Seurat’s SCTransform method (version 2 [80]), utilizing the following functions (with default parameters): ‘NormalizeData’, ‘FindVariableFeatures’, ‘ScaleData’, ‘SCTransform’, ‘RunPCA’, ‘RunUMAP’. The PCA was performed on the variable features. The UMAP was constructed from the first 30 principal components.

#### Clustering and cell type identification

For each compartment, nearest neighbor identification and unsupervised clustering were also performed using functions from the Seurat package (default parameters): ‘FindNeighbors’ and ‘FindClusters’. We examined resolutions 0.1 through 0.9, with a step size of 0.1. For each compartment, cell types were identified by (1) selecting a clustering resolution which recapitulated known biological priors and (2) annotating clusters based on the expression of known markers and signature scores. Then, clusters were merged into biologically relevant groupings (e.g. two cell clusters identified by unsupervised clustering which express *CD8A* and *CD3E* labelled as “CD8+ T cell”; referred to as “general cell type” in Supplementary Fig. 7b). General cell type markers can be found in Supplementary Fig. 7c.

For immune cell subsets, further sub-clustering was done on the B cells, myeloid cells, and T/NK cells to identify specific immune cell subtypes. When differentiating between very similar immune cell subsets, sample-level differences in phenotype oftentimes dominated (e.g. FOXP3+ T regulatory cells clustering with other T cells from the same sample rather than clustering together). To mitigate these sample-level differences which arise when trying to distinguish between similar cells, we integrated the cell type subset using CCA integration described in [81] and implemented in the Seurat package, utilizing the functions: ‘SelectIntegrationFeatures’ (3000 features), ‘PrepSCTIntegration’, ‘FindIntegrationAnchors’ (first 15 principal components), and ‘IntegrateData’ (considering 15 neighbors when weighting anchors). Then, for each subset, we re-ran the same workflow as described above, identifying finer immune cell subsets (referred to as “specific cell type”). Specific cell type markers can be found in Supplementary Fig. 7d.

For additional clarification about the exhaustion phenotype in the CD8+ T cell subset, CD8+ T cells were additionally scored with signatures derived from tumor subsets in Miller, Sen et al. 2019 [47] (Supplementary Fig. 7e).

#### Per-cell signature scoring

All per-cell signature scoring in the scRNA-seq analysis was done using AUCell (v1.20.2) [82], a rank-based method for assigning scores using gene signatures. A subset of the signatures utilized in the bulk RNA-seq analysis were used in the scRNA-seq signature scoring. Additionally, the SPP1+ macrophage signature from supplementary tables in Wei et al. 2021 [43] was downloaded.

#### Calculation of sample-level hypoxia score

The sample-level hypoxia score was calculated from the AUCell signature scores, utilizing the ‘HALLMARK_HYPOXIA’ signature from MSigDB. We noted that the signature score was significantly different between the tumor, stromal, and immune compartments, with stromal cells expressing the highest levels of these signatures and immune cells expressing the lowest. To decouple the sample-level hypoxia score from the relative proportion of stromal, tumor, and immune cells, we first normalized the hypoxia score for every cell within each compartment, effectively z-scoring the distribution of hypoxia scores per cell for immune, tumor, and stromal cells separately. For compartment-level hypoxia scores, we calculated the mean hypoxia score for each compartment and sample. For the sample-level hypoxia scores, we calculated the mean hypoxia score using the normalized hypoxia score for each sample across all cells in that sample.

#### Pseudobulk analysis and modeling sample-level hypoxia

Pseudobulk samples were creating by aggregating the counts for each gene across all cells for each sample and cell type using functions documented in the custom package Rsc (v0.0.900, hosted at github.com/ amyh25/Rsc). For the pseudobulk analysis, since we wanted to enrich for higher quality cells, we only included immune and stromal cells with greater than 1000 unique features. For tumor cells, we did not exclude any cells, since we could not confidently exclude cells with fewer features as simply lower quality cells due to the complexities of identifying and analyzing tumor cells in scRNA-seq. For the SCTransform normalized Seurat objects, the SCT transformed counts were taken instead of the RNA counts. These functions remove cells which are outliers by median absolute deviation (MAD) and consider only genes with greater than 10 total counts for each pseudobulk, using functions from scater (v1.26.1) and Matrix.utils (v0.9.8). DESeq2 (v1.38.3) was used to normalize the pseudobulk.

To understand how gene expression in specific cell types related to the overall sample-level of hypoxia, we constructed linear models to predict sample-level hypoxia from the expression of genes for each cell type (model formulation: hypoxia score ∼ gene expression). For the gene expression, we utilized the log-normalized counts per million (log(CPM + 1)). To enrich for the most relevant gene changes in each pseudobulk, we only considered genes with log2 fold change greater than 0.1. We then used the estimated effect of the hypoxia for downstream signature scoring, in a similar analysis to the bulk RNA-seq (see section on pre-ranked gene set enrichment analysis in bulk RNA-seq analysis for details).

#### Receptor-ligand analysis

Receptor-ligand analysis was performed using the algorithm and database from CellPhoneDB [45]. Similarly to the pseudobulk analysis, we only included immune and stromal cells with greater than 1000 unique features. Metadata and the input counts matrix (pulled from the RNA “slot” of each Seurat object) was merged into a table for each sample. Then, the Cell-PhoneDB analysis was run from command line using cellphonedb method statistical_analysis. Downstream processing and visualization of the Cell-PhoneDB output was done in R using functions from the tidyverse set of packages (v2.0.0).

### Additional scRNA-seq analysis of clinical datasets

Two addition scRNA-seq cohorts of metastatic melanoma were considered for replication of the hypoxia stratification analysis. We identified the scRNA-seq dataset published in Jerby-Arnon et al. 2018 [44] and the scRNA-seq dataset published in Sade-Feldman et al. 2018 [11] as suitable datasets to reproduce the analysis. While the sequencing modality of these datasets (SmartSeq2) were different from the Yang et al. 2024 scRNA-seq cohort, both datasets contained pre-treatment samples from patients with metastatic melanoma. Note that compared to the Yang scRNA-seq cohort, these datasets are more limited by the number of cells detected. In particular, this made it infeasible to replicate the stromal cell receptor-ligand analyses, so instead analysis was focused on the immune compartment results. The counts matrices, cell- and sample-level metadata of the external single cell RNA-sequencing cohort was downloaded from the supplementary material of both publications. From the counts matrices, samples were processed using the standard Seurat workflow up through dimensionality reduction. In the Jerby-Arnon et al. 2018 dataset, we note the following about the analysis: cell types were not re-annotated, although they were re-labelled to match with the cell type labels from the scRNA-seq analysis. Only metastatic samples from the “Untreated” treatment group were considered for the downstream hypoxia analysis. The analysis of the effects of hypoxia were conducted as described above also in the external scRNA-seq cohort. In the Sade-Feldman et al. 2018 dataset, we note the following about the analysis: given that this dataset only contained immune cells, the hypoxia stratification was conducted with only immune cells, and thus the results may be different from the other two datasets. Cell types were re-annotated post-clustering.

### Expression of hypoxia-related genes from B16 tumors

FPKM expression matrices from bulk RNA-sequencing of B16 tumors was downloaded from GEO (accession number: GSE96972) [46]. In this study, B16 tumor cells were seeded on extracellular matrix and treated *in vitro* with siRNA in two types of media. Expression of hypoxia-related genes were taken from conditions treated with non-targeting siRNA (siNT). The gene encoding succinate dehydrogenase complex flavoprotein subunit A (*Sdha*) was chosen as a representative house-keeping gene.

### Expression of hypoxia-related genes from B16 TME

TPM expression matrices from scRNA-sequencing of B16 tumors (the murine melanoma atlas) were downloaded from https://www.ebi.ac.uk/gxa/sc/experiments/E-EHCA-2/downloads. Cluster annotations, as described in Davidson et al. 2020 [83], were also downloaded. Only clusters which were present in the tumor were considered; lymph node clusters were discarded. The following clusters were merged for clarity of visualization: conventional dendritic cells (DCs), migratory DCs, plasmacytoid DCs were all labelled “DCs”; cancer-associated fibroblast clusters 1 through 3 were all labelled “fibroblasts”; gamma delta T cells/mucosal-associated invariant T cells along with tumor T cells were all labelled “T cells,” macrophages and monocytes were labelled as “TAMs” for tumor-associated macrophages and monocytes. Gene expression was then visualized as log_10_(TPM + 1).

#### Visualizations

Visualizations were created with the R packages ggplot2 (v3.4.2), cowplot (v1.1.1), ggpmisc (v0.5.5), ggpubr (v0.6.0), scattermore (v1.2), ComplexHeatmap (v2.6.2), and survminer (v0.4.9). The Adobe Illustrator software was used to lay out the visualizations and harmonize the visual style.

#### Tumor cell lines

B16-OVA expressing cell lines were generated from B16.F10 as described previously [47]. B16.F10 and B16-OVA cells were cultured in Dulbecco’s modified Eagle’s medium (DMEM, Gibco) with 5% fetal bovine serum (FBS, Gemini Bio-Products) and 1% Penicillin-Streptomycin (Life Technologies) at 37°C with 5% CO_2_ and passaged 3-5 times prior to implantation. Culture media was supplemented with 2 ug/ml puromycin for B16-OVA cells. Cell lines were tested intermittently for mycoplasma contamination using the LookOut Mycoplasma PCR detection kit.

#### Mice and tumor implantation

Female 6-8 week old C57BL/6J mice (Catalog #000664) were purchased from the Jackson Laboratory. All mice were maintained in specific pathogen-free facilities at Harvard Medical School under standard housing, husbandry, and diet conditions according to Institutional Animal Care and Use Committee and NIH guidelines. All experimental procedures performed were approved by the Institutional Animal Care and Use Committee at Harvard Medical School.

Mice were implanted subcutaneously with 3 *×* 10^5^ B16.F10 or 3 *×* 10^5^ B16-OVA in the flank. Once the average tumor size across cages reached *>*100 mm^3^, cages were normalized, and dosing schedules began. In vivo antibodies were sourced from BioXCell. All blocking antibodies and isotype controls were administered via intraperitoneal (i.p.) injection in cold DPBS^−*/*−^ in 200µl volumes. 200µg anti-PD-1 (clone 29 F.1A12) or 200µg isotype-matched control antibody (2A3, Rat IgG2a) were administered. PT2399 (MedChem Express) was administered by oral gavage, twice daily, between days 11-25 at 30mg/kg (0.75mg) doses. To prepare the dosing solution, PT2399 was dissolved in 10% ethanol, 30% polyethylene glycol 400 (Sigma-Aldrich) and further diluted in 0.5% (hydroxypropyl)methyl cellulose (HPMC) solution (Sigma-Aldrich). The dosing solution was sonicated for 20 minutes prior to administration. Mice were monitored for tumor growth starting on day 7 or 8 with tumor volume calculated as the volume of an ellipsoid (0.5 *× D × d*^2^) where *D* refers to the largest diameter and *d* refers to the smaller diameter of the tumor. Mice were sacrificed on the days specified for tissue dissection or when they reached humane endpoint: 2000mm^3^ or significant ulceration. All experiments were repeated with reproducible results.

#### Flow cytometry

Mice were euthanized and the tumors were manually dissected. Tumors were mechanically minced and digested with type I collagenase (2mg/ml, Worthington Chemical) for 20 minutes at 37°C. Tumor infiltrating lymphocytes were enriched using a 70%/40% Percoll gradient with centrifugation at 2000 rpm at 400 x g for 20 minutes at 20°C without acceleration or brake.

Single cell suspensions of lymphocytes from tumors were pre-treated with anti-CD16/CD32 Mouse Fc block (BD Biosciences 553142, 1:100) following by staining with fluorescently labelled antibodies against: CD45.2 BUV395 (clone 104, BD 564616, 1:200), Ki-67 (clone B56, BD 561284, 1:300), CD3e BUV737 (145-2C11, BD 612771, 1:100), CD4 BUV496 (clone GK1.5, BD 612952, 1:100), CD8a BUV805 (clone 53-6.7, BD 612898, 1:100), Granzyme B Pacific Blue (clone GB11, Biolegend 515407, 1:100), TIM-3 BV711 (clone RMT3-23, Biolegend 119727, 1:200), Foxp3 FITC (clone FJK-16s, ThermoFisher 11-5773-82, 1:200), PD-1 PE-Cy7 (clone RMP1-30, Biolegend 109110, 1:200), and CD25 PE Cy5 (clone PC61, Biolegend 102010, 1:200). In a separate flow cytometry panel, cells were stained with fluorescently labelled antibodies against CD45.2 BUV605 (clone 104, Biolegend 109841, 1:200), CD31 BUV563 (clone 390, BD 741262, 1:100), Vimentin AlexaFluor488 (clone O91D3 Biolegend 677809, 1:200), and HIF1a (clone 241812, R&D IC1935R, 1:100). Dead cells were excluded using Zombie NIR Fixable Viability kit (Biolegend 423106, 1:600) or Zombie Aqua (Biolegend, 423101, 1:500). Intracellular staining against Ki-67, granzyme B, and Foxp3 was performed using the eBioscience Foxp3 transcription factor fixation/permeabilization kit per manufacturer’s instructions (Thermo Fisher Scientific). Data were acquired on a FACS Symphony A5 (BD Biosciences) using BD FACS-Diva v9.0 (BD Biosciences) or the Aurora Spectral Analyser (Cytek Biosciences). Analysis was performed with FlowJo (v10.8.1, BD Biosciences). Autofluorescence extraction was applied to experimental spectral data using unstained tissues subjected to identical processing conditions. Cell counts were obtained using the Cytek’s built-in counting software or CountBright− Absolute Counting Beads (Invitrogen) per manufacturer’s instructions. All experiments were repeated with reproducible results.

#### Statistical analysis

For all transcriptomic analyses, statistical analyses were performed in R (v4.2.1 “Funny Looking Kid”). Reported p-values represent nominal p-values unless otherwise specified. Unless otherwise stated, we used the Mann-Whitney U-Test for any comparison between two groups of continuous clinical or molecular features and Fisher’s Exact Test for the association of binary variables. All statistical tests performed were two-sided unless otherwise stated.

For flow cytometry analyses, statistical analysis was performed using GraphPad Prism 9 (v9.5.1). For comparison of means between two groups, an unpaired Student’s t-test was utilized. Analysis for multi-group multivariate statistics comparing multiple means was performed using a two-way ordinary ANOVA (95% CI), with post-hoc analysis of Tukey’s multiple comparisons test for comparisons of all means within the test group for multiple-comparison correction. P values *<*0.05 were considered statistically significant.

## Supplementary Figures

**Supplementary Figure 1.**
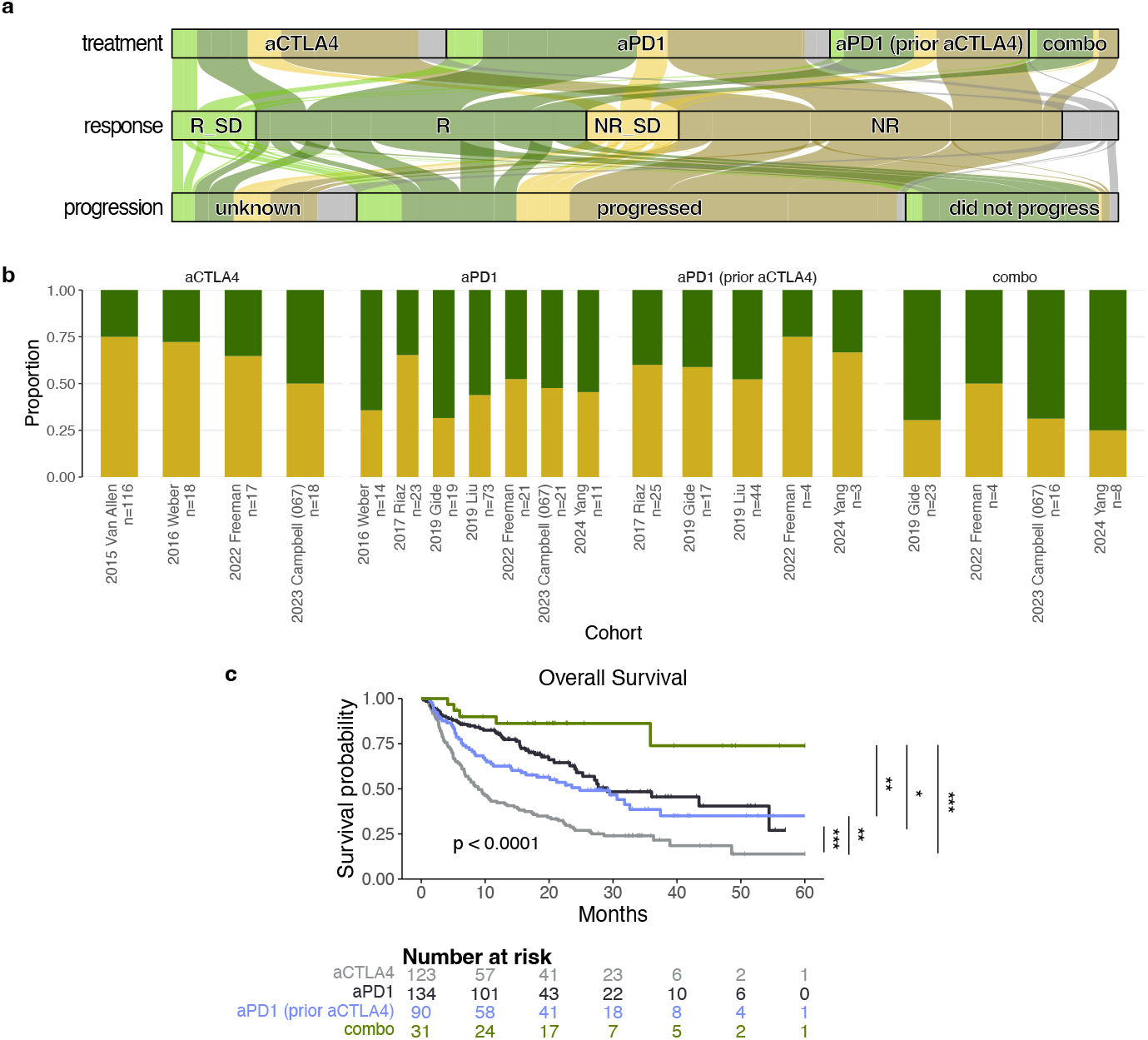
Additional clinical details about the bulk sequencing meta-cohort. **a**. Subset of samples with response annotations and bulk RNA-sequencing, shown in a Sankey diagram. **b**. Overall survival rates stratified by treatment group. Treatment groups are significantly different (Log-rank *p <* 0.0001). Significance of pairwise log-rank also shown. **c**. Response rates by treatment group and immune state in each cohort follows trends for the aggregated meta-cohort.

**Supplementary Figure 2.**
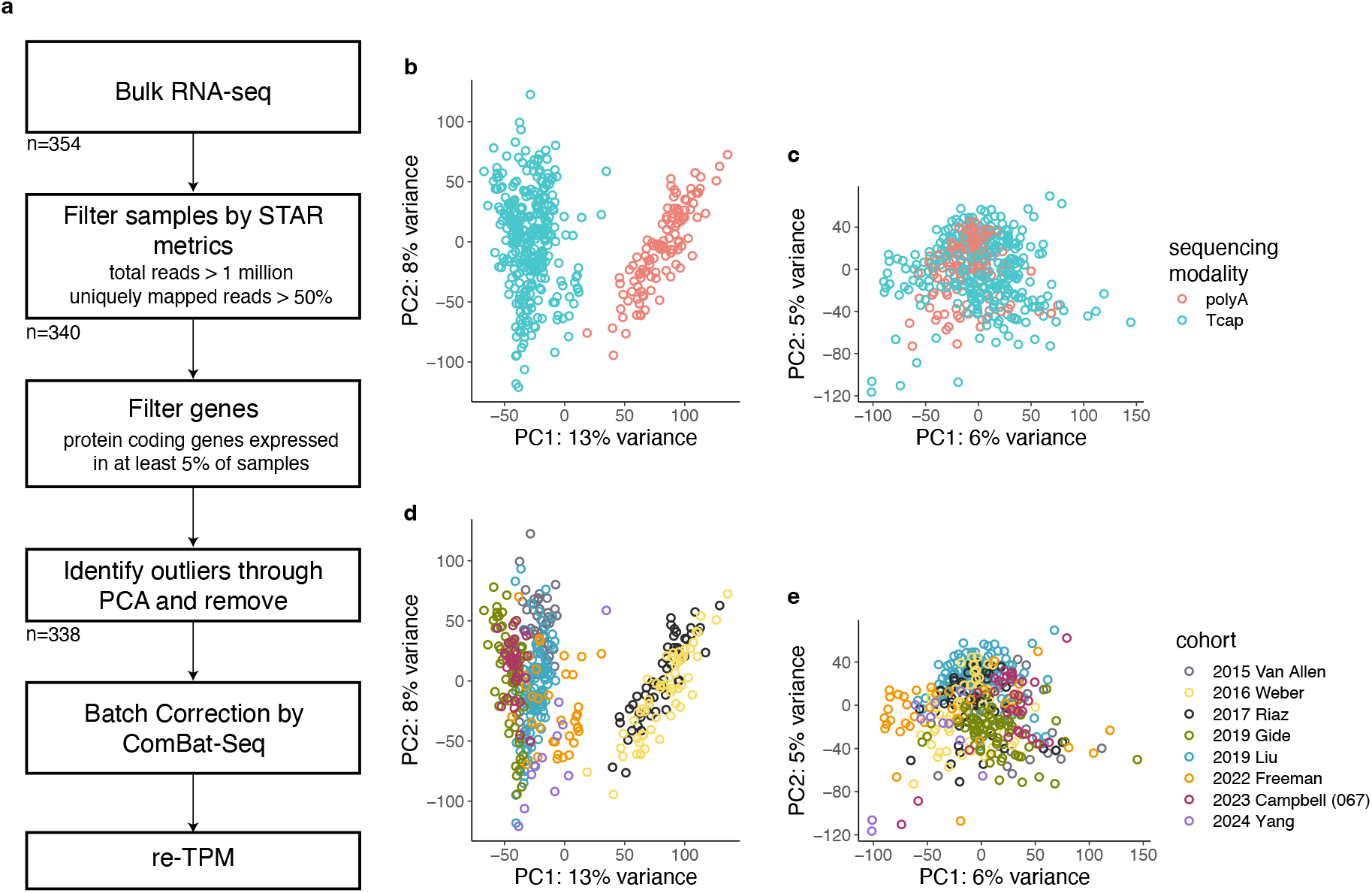
Bulk RNA-seq processing. **a**. Graphic depicting bulk RNA-seq processing workflow. **b**. Principal component analysis (PCA) of bulk RNA-seq samples before batch correction. Colored by sequencing modality. **c**. PCA of bulk RNA-seq samples after batch correction. Colored by sequencing modality. **d**. Same as **b**., but colored by cohort. **e**. Same as **c**., but colored by sequencing modality.

**Supplementary Figure 3.**
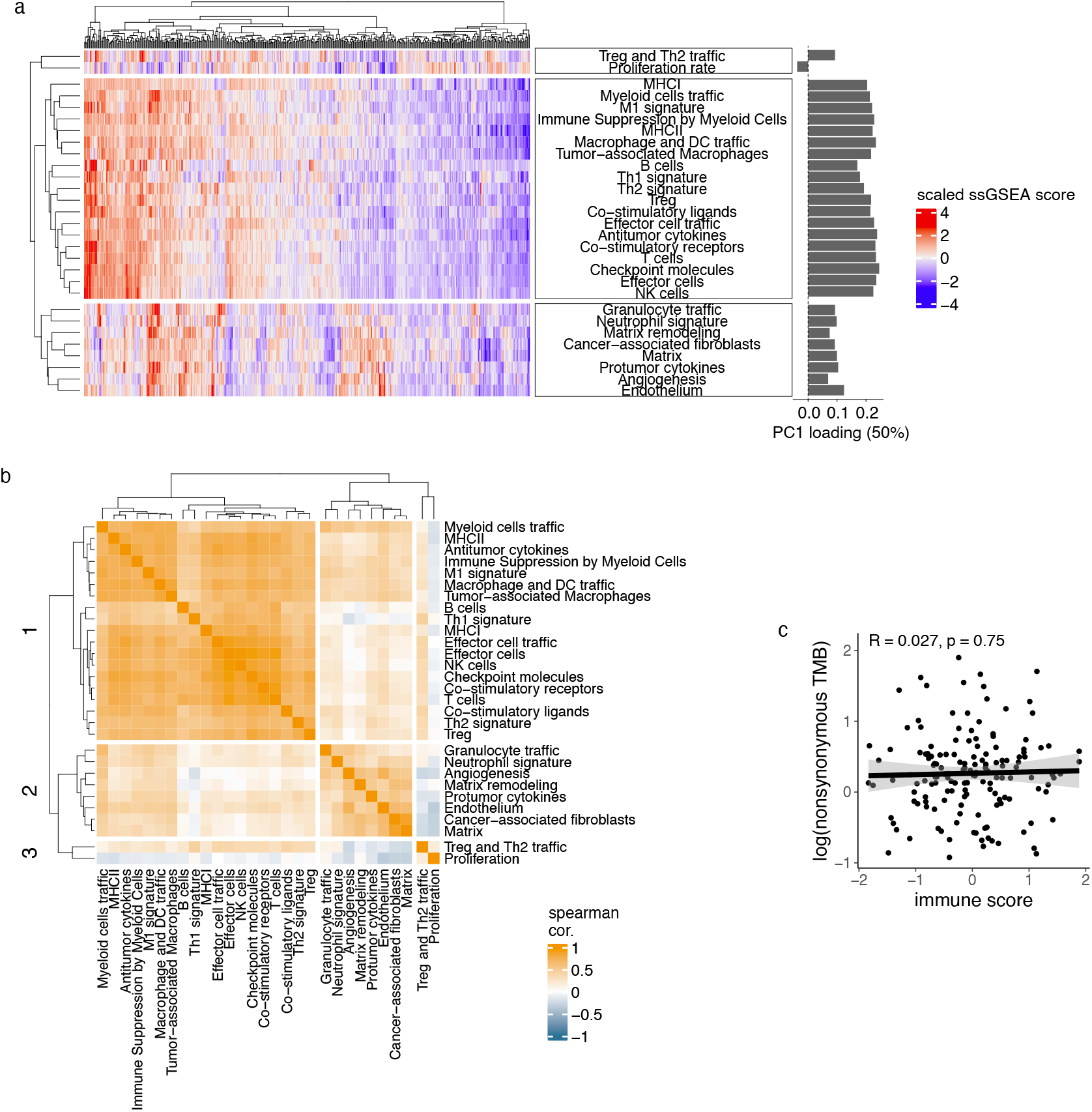
Characteristics of cell type signatures scores and immune high versus low categories. **a**. Heatmap with samples hierarchically clustered based on Euclidean distance between samples’ cell type signature scores (for all signatures from Bagaev et al. 2021 [25]). Loadings from principle component analysis for each signature are shown to the right of the heatmap. The first principle component explains 50% of the variance between the samples. **b**. Co-correlation heatmap for all signatures from Bagaev et al. 2021 [25]. Correlations between each group are Spearman’s rho *ρ*. For all comparisons between the 19 immune cell type signatures, the correlation is between 0.68 *±* 0.11 (mean *±* standard deviation). **c**. Scatterplot showing no relationship between immune score and tumor mutational burden (TMB).

**Supplementary Figure 4.**
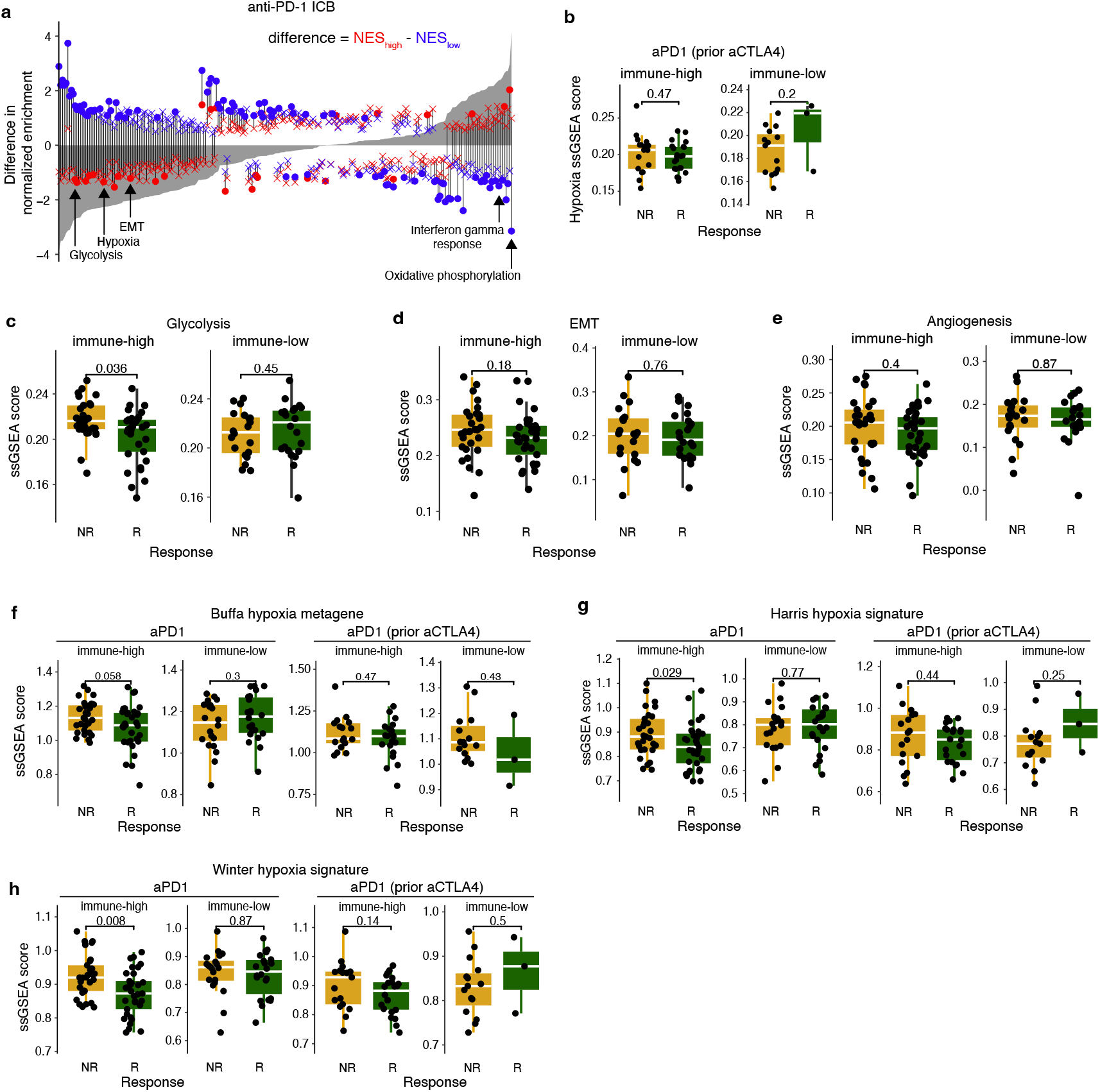
Hypoxia-related signatures and response to anti-PD-1 considering immune stratification. **a**. Difference in normalized enrichment score for response to anti-PD-1 between immune-high and immune-low for each signature (grey profile). Signatures are ordered by difference between immune-high and immune-low. Ends of the vertical bar show the enrichment of the signature in immune-high (red) and immune-low (blue). Specific pathways are highlighted by arrows (from left to right): hypoxia, interferon gamma response, oxidative phosphorylation. **b**. Difference between ssGSEA scores of labelled gene sets in responders and non-responders, split by immune high and low, in patients treated with anti-PD-1 without prior anti-CTLA-4. **c-e**. Difference between ssGSEA scores for the labelled signatures in responders and non-responders, split by immune high and low, in patients treated with anti-PD-1. From left to right: glycolysis, epithelial mesenchymal transition (EMT), and angiogenesis. **f-h**. Difference between three additional hypoxia signature ssGSEA scores (**f**. [30], **g**. [31], **h**. [29]) in responders and non-responders, split by immune high and low, in patients treated with anti-PD-1 with and without prior anti-CTLA-4. Signatures were access through the MSigDB database (see Methods) [26].

**Supplementary Figure 5.**
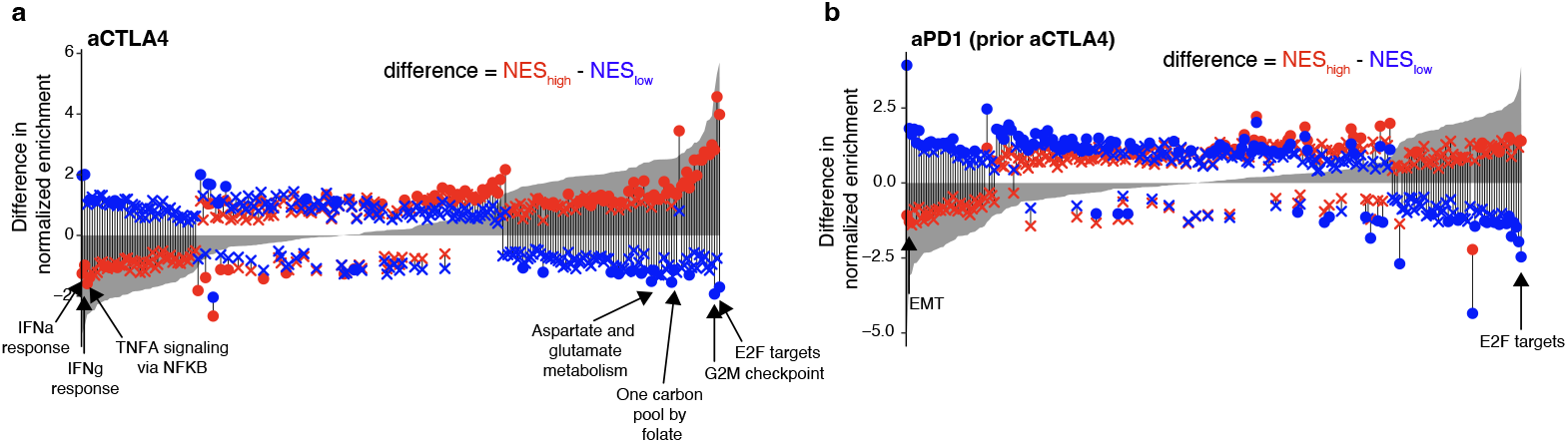
Differences in GSEA results between immune-high and immune-low for additional treatment regimens. **a-b**. Difference in normalized enrichment score for response to **a**. anti-CTLA-4 and **b**. anti-PD-1 with prior anti-CTLA-4 between immune-high and immune-low for each signature (grey profile). Signatures are ordered by difference between immune-high and immune-low. Ends of the vertical bar show the enrichment of the signature in immune-high (red) and immune-low (blue). Specific pathways are highlighted by arrows.

**Supplementary Figure 6.**
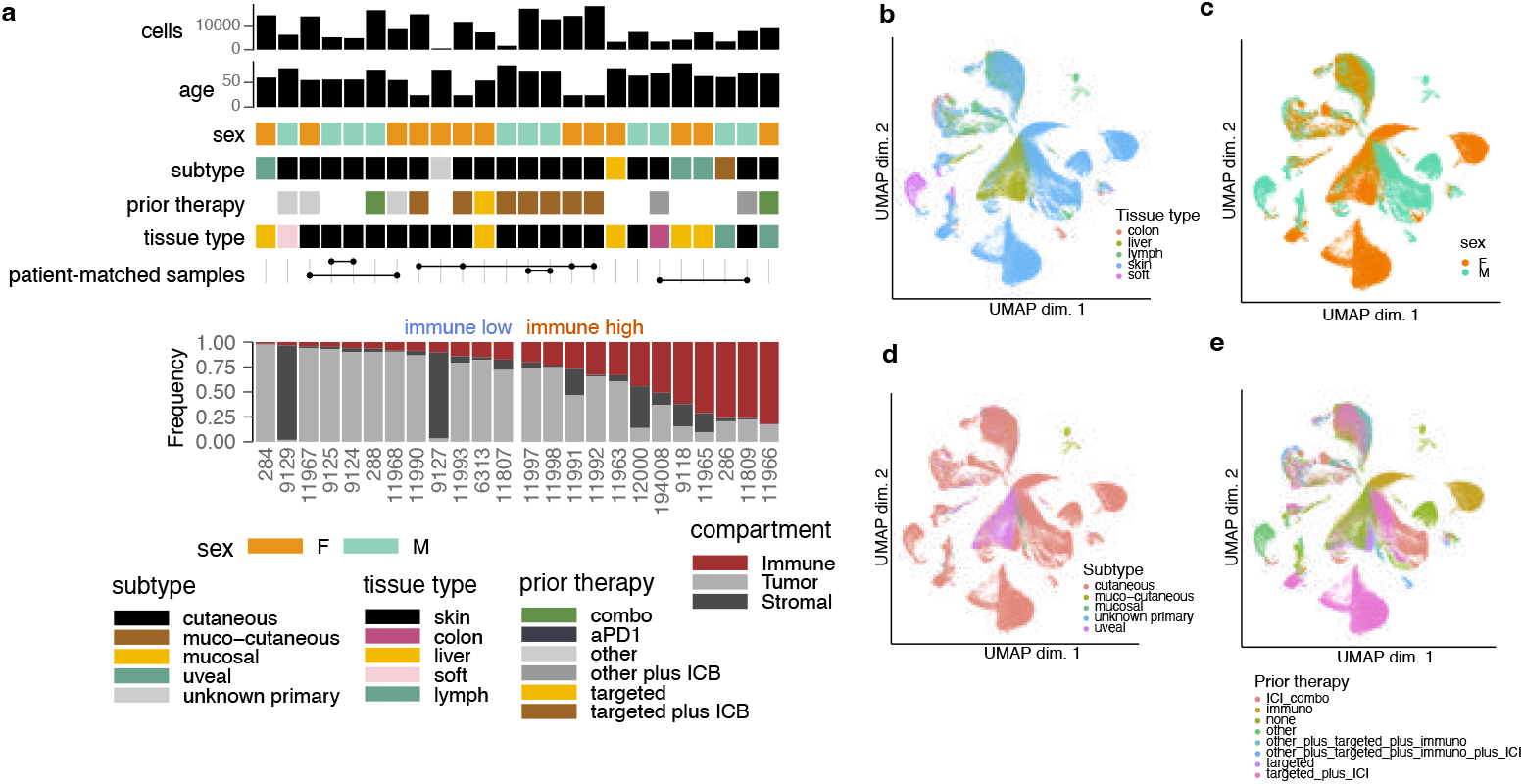
Clinical features of single-cell cohort. **a**. Clinical features of the single-cell cohort. From the top row to the bottom row: the number of cells, age of the patient, sex of the patient, melanoma subtype (if known), prior therapy, tissue type of biopsy, samples from the same patient, proportion of immune cells and immune state. **b-e**. Clinical features (**b**. tissue type, **c**. sex, **d**. subtype, **e**. prior therapy) projected onto UMAP.

**Supplementary Figure 7.**
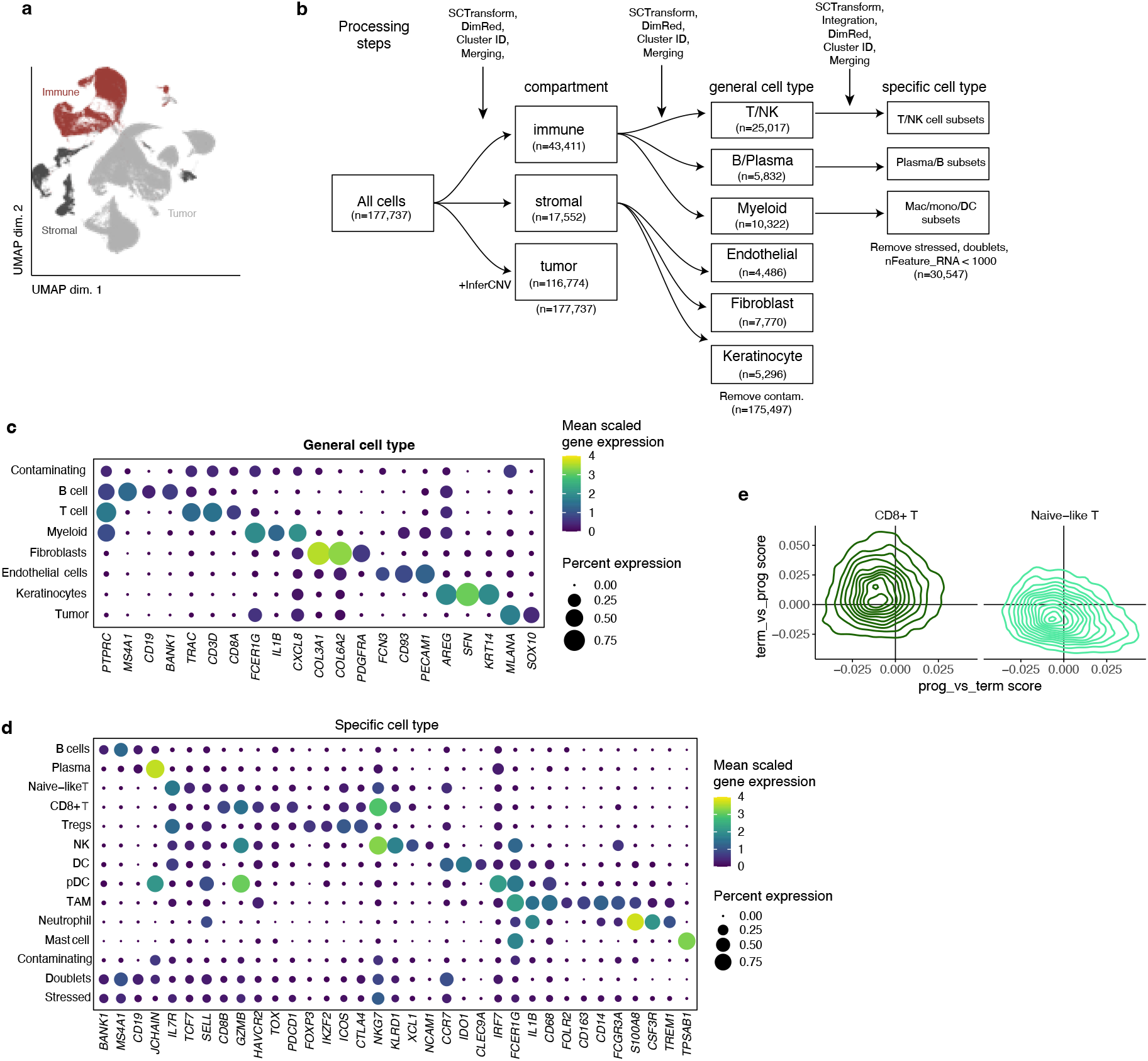
Clustering and cell identification in single cell RNA-seq cohort. **a**. UMAP with compartment-level clustering annotations. **b**. Diagram detailing sub-clustering workflow for single-cell RNA-seq data, pre-treatment subset. **c**. Markers of general cell types shown in dot plot. **d**. Markers of specific cell types shown in dot plot. **e**. CD8+ T and Naive T cell clusters scoring with progenitor versus terminal CD8+ T exhaustion signature from tumor data in Miller et al. 2019 [47]. In dot plots, dot size indicates percentage expressed, color indicates mean expression.

**Supplementary Figure 8.**
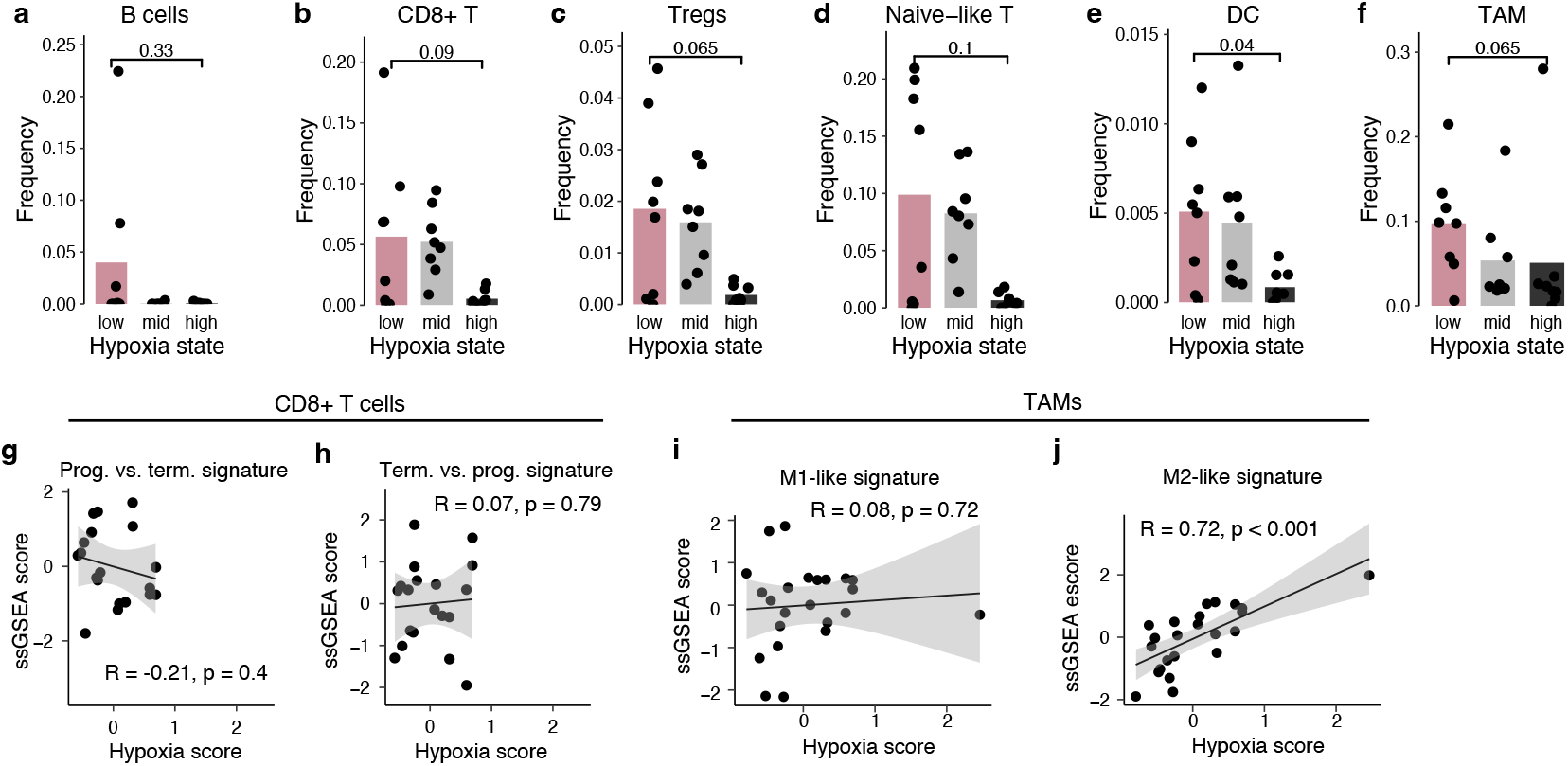
Additional associations with hypoxia in the scRNA-seq cohort. **a-f**. Proportion of total immune cells for each sample by the listed cell type, varying across hypoxia state (low, mid, high). **g-h**. The relationship between the normalized ssGSEA score of the progenitor versus terminally exhausted CD8+ tumor-infiltrating lymphocyte signature (TIL) from Miller et al. 2018 [47] in the CD8+ T cell pseudobulks and the sample-level hypoxia score. **i-j**. The relationship between the normalized ssGSEA score of M1-like and M2-like gene signatures in TAM pseudobulks and the sample-level hypoxia score.

**Supplementary Figure 9.**
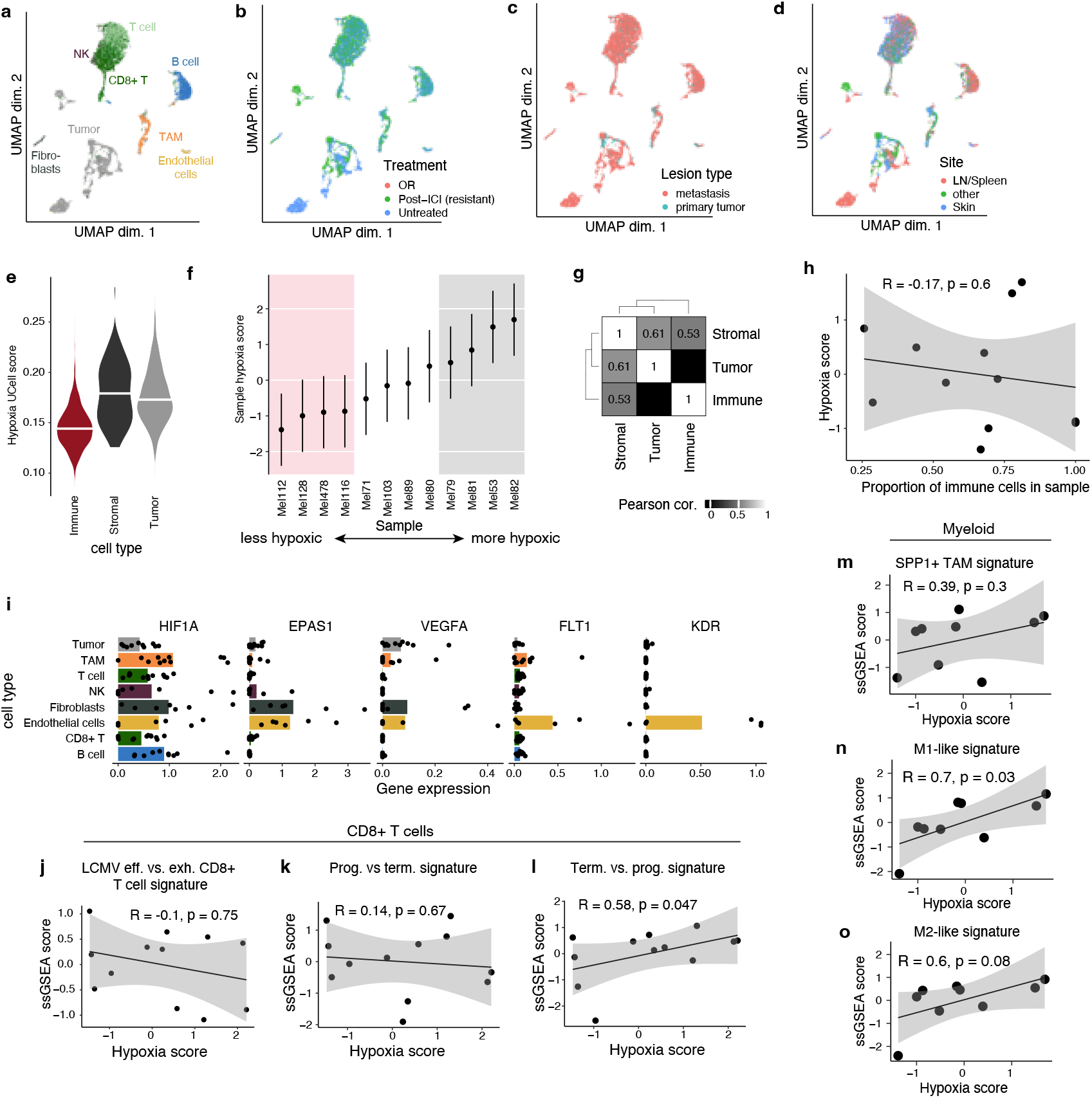
Replication of hypoxia analysis in Jerby-Arnon et al. 2018. **a**. UMAP with cell type annotations of all cells from scRNA-seq published in Jerby-Arnon et al. 2018 [44]. **b-d**. UMAP with clinical annotations (Treatment group, lesion type) projected. Only metastases which were untreated were considered in the subsequent analysis. **e**. Hallmark Hypoxia score in each general cell type compartment. **f**. Sample-level hypoxia score for all “Untreated” samples. **g**. Co-correlation between hypoxia score in each general compartment shows correlation between tumor and stromal compartments (Pearson’s *r* = 0.61). **h**. Relationship between sample-level hypoxia and proportion of immune cells. **i**. Expression of key hypoxia-related transcription factors (*EPAS1* (HIF-2*α*), *HIF1A*) and downstream angiogenesis-mediating genes (*FLT1, KDR, VEGFA*) across cell types. **j**. Normalized ssGSEA score of an LCMV effector versus exhausted CD8+ T cell signature (GSE41867_DAY15_EFFECTOR_VS_DAY30_EXHAUSTED_CD8_-TCELL_LCMV_CLONE13) in CD8+ T cell pseudobulks correlated with sample-level hypoxia score. **k-l**. The relationship between the normalized ssGSEA score of the progenitor versus terminally exhausted CD8+ tumor-infiltrating lymphocyte signature (TIL) from Miller et al. 2018 [47] in the CD8+ T cell pseudobulks and the sample-level hypoxia score. **m-o**. The relationship between the normalized ssGSEA score of M1-like and M2-like gene signatures as well as the SPP1+ TAM signature from Wei et al. 2021 [43] in TAM pseudobulks and the sample-level hypoxia score.

**Supplementary Figure 10.**
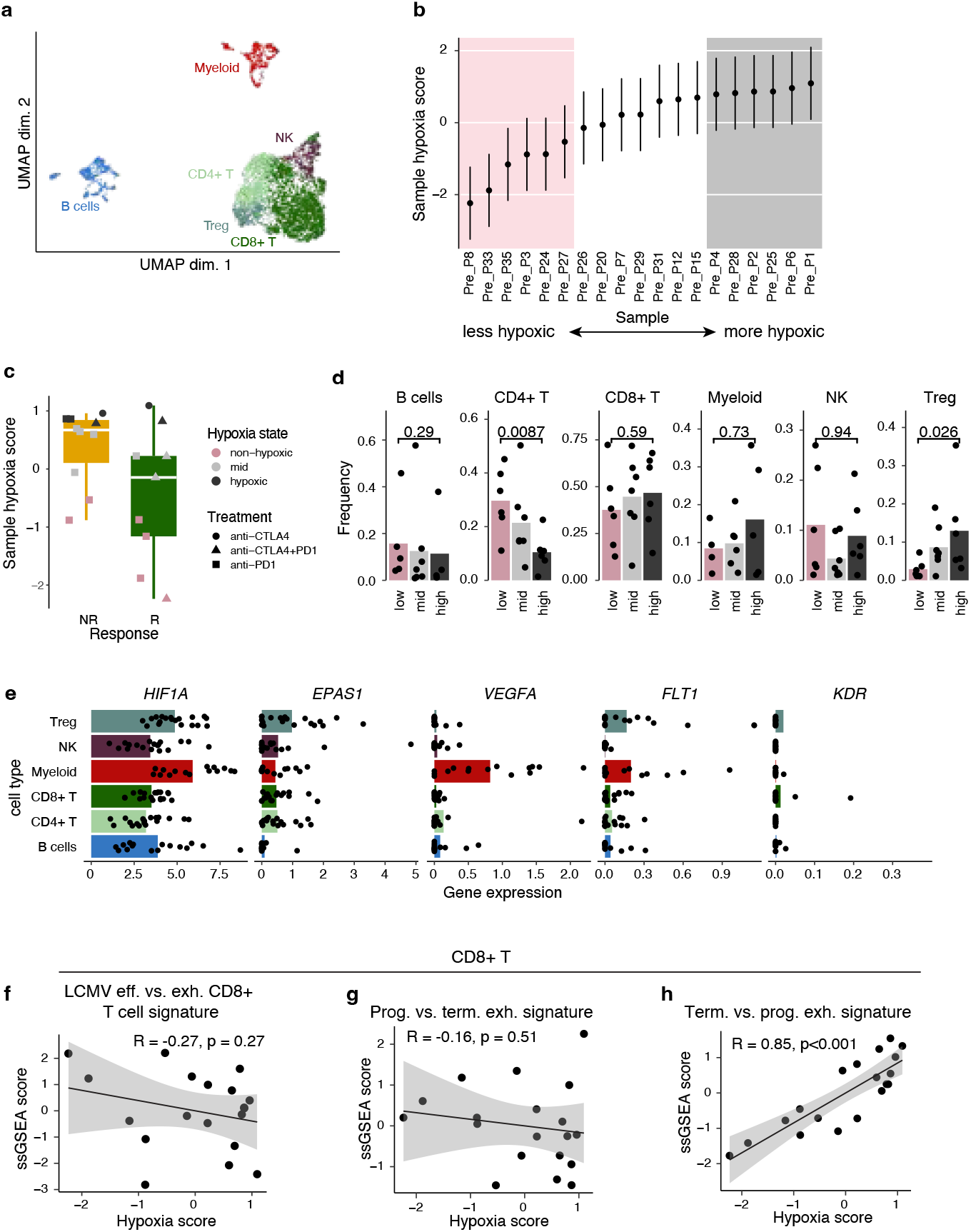
Replication of hypoxia analysis in Sade-Feldman et al. 2018. **a**. UMAP with cell type annotations of all cells from scRNA-seq published in Sade-Feldman et al. 2018 [11]. **b**. Sample-level hypoxia score for all pre-treatment samples. **c**. Sample-level hypoxia score in responders and non-responders for all pre-treatment samples. Treatment regimen and hypoxia state are encoded in shape and color. **d**. Proportion of total immune cells for each sample by the listed cell type, varying across hypoxia state (low, mid, high). **e**. Expression of key hypoxia-related transcription factors (*EPAS1* (HIF-2*α*), *HIF1A*) and downstream angiogenesis-mediating genes (*FLT1, KDR, VEGFA*) across cell types present in the dataset. **f**. Normalized ssGSEA score of an LCMV effector versus exhausted CD8+ T cell signature (GSE41867_DAY15_EFFECTOR_VS_DAY30_EXHAUSTED_CD8_-TCELL_LCMV_CLONE13) in CD8+ T cell pseudobulks correlated with sample-level hypoxia score. **f-h**. The relationship between the normalized ssGSEA score of the progenitor versus terminally exhausted CD8+ tumor-infiltrating lymphocyte signature (TIL) from Miller et al. 2018 [47] in the CD8+ T cell pseudobulks and the sample-level hypoxia score.

**Supplementary Figure 11.**
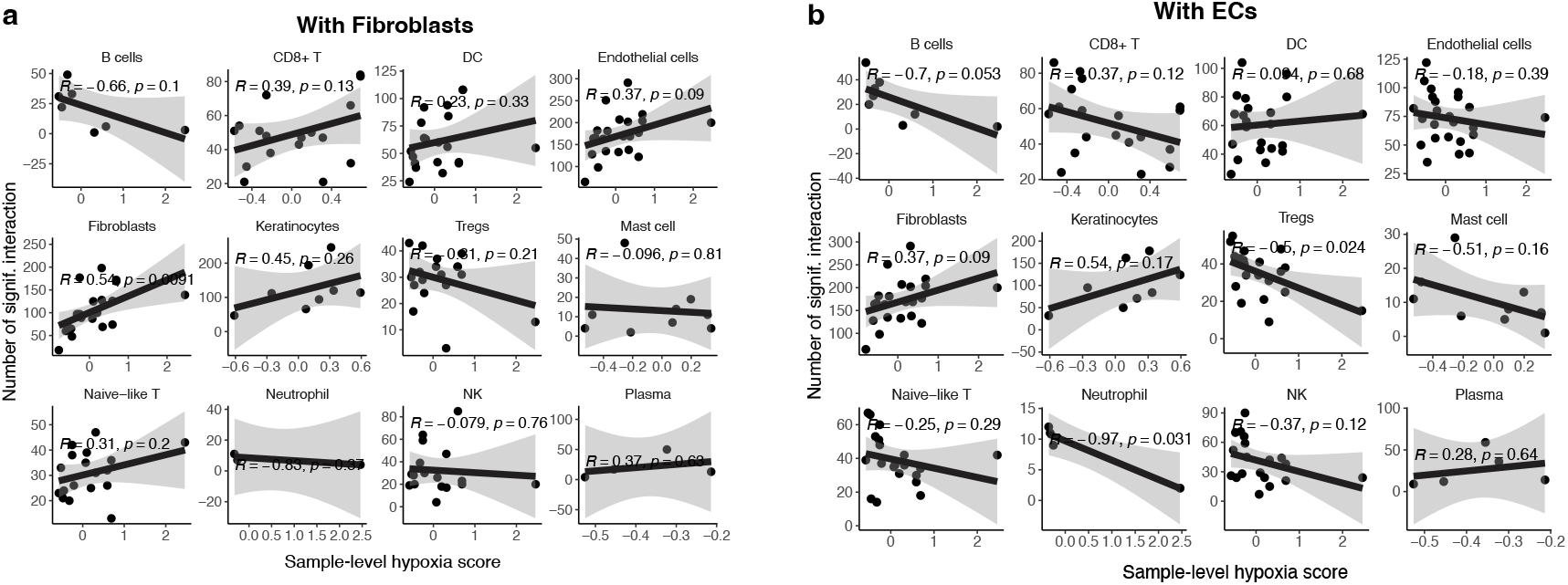
Stromal interactions in the tumor microenvironment. **a-b**. The relationship between the number of stromal interactions estimated by the CellPhoneDB statistical analysis method and the sample-level hypoxia score for **a**. fibroblasts and **b**. endothelial cells (ECs). Correlations shown are Pearson correlations.

**Supplementary Figure 12.**
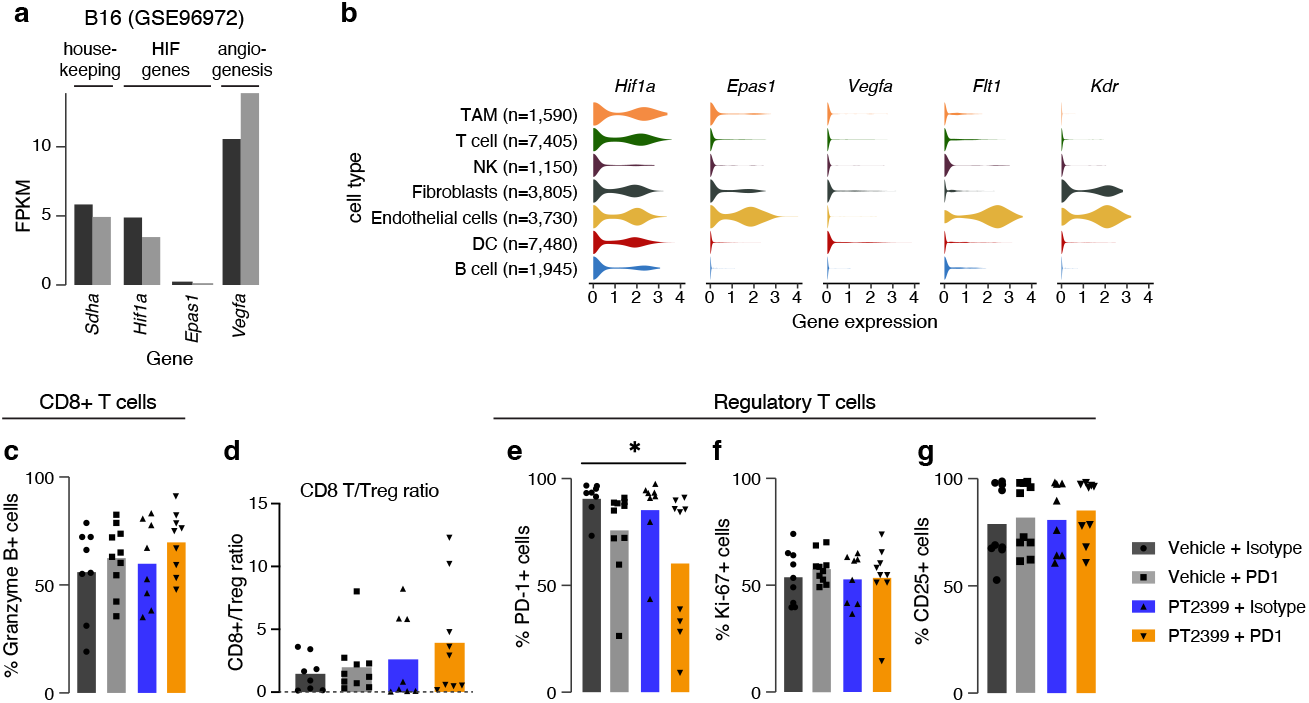
Characterization of T cells in B16-OVA tumors from mice treated with combined PT2399/anti-PD-1 therapy. **a**. Gene expression (FPKM) of hypoxia-related genes in B16 tumors. Left bar (black) represents expression in serum free media (SFM) and right bar (grey) represents expression in tumors in tissue culture media (TCM). Data from Murgai et al. 2017 (GSE96972) [46], which describes tumor cells as seeded on extracellular matrix and treated with siRNA. Only data from tumor cells treated with non-targeting RNA (siNT) are shown. **b**. Gene expression (log_10_(TPM + 1)) of hypoxia-related genes for different cell types in the B16 tumor microenvironment (TME). Data from the murine melanoma atlas as described in Davidson et al. 2020 [83]. **c-g**. Flow cytometry characterization of T cells isolated from B16-OVA tumors treated with isotype control, anti-PD-1, PT2399, or the combination. Data showing two independent experiments combined (n=4-5 per group). **c**. Percentage of GzmB+ T cells of CD8+ T cells. **d**. Ratio CD8+ T cells to regulatory T cells in B16-OVA tumors treated with isotype control, anti-PD-1, PT2399, or the combination. **e**. Percentage of PD-1+ T cells of Tregs. **f**. Percentage of Ki-67+ T cells of Tregs. **g**. Percentage of CD25+ T cells of Tregs. Significance levels are denoted as \**p <* 0.05, \*\**p <* 0.01, \*\*\**p <* 0.001.

**Supplementary Figure 13.**
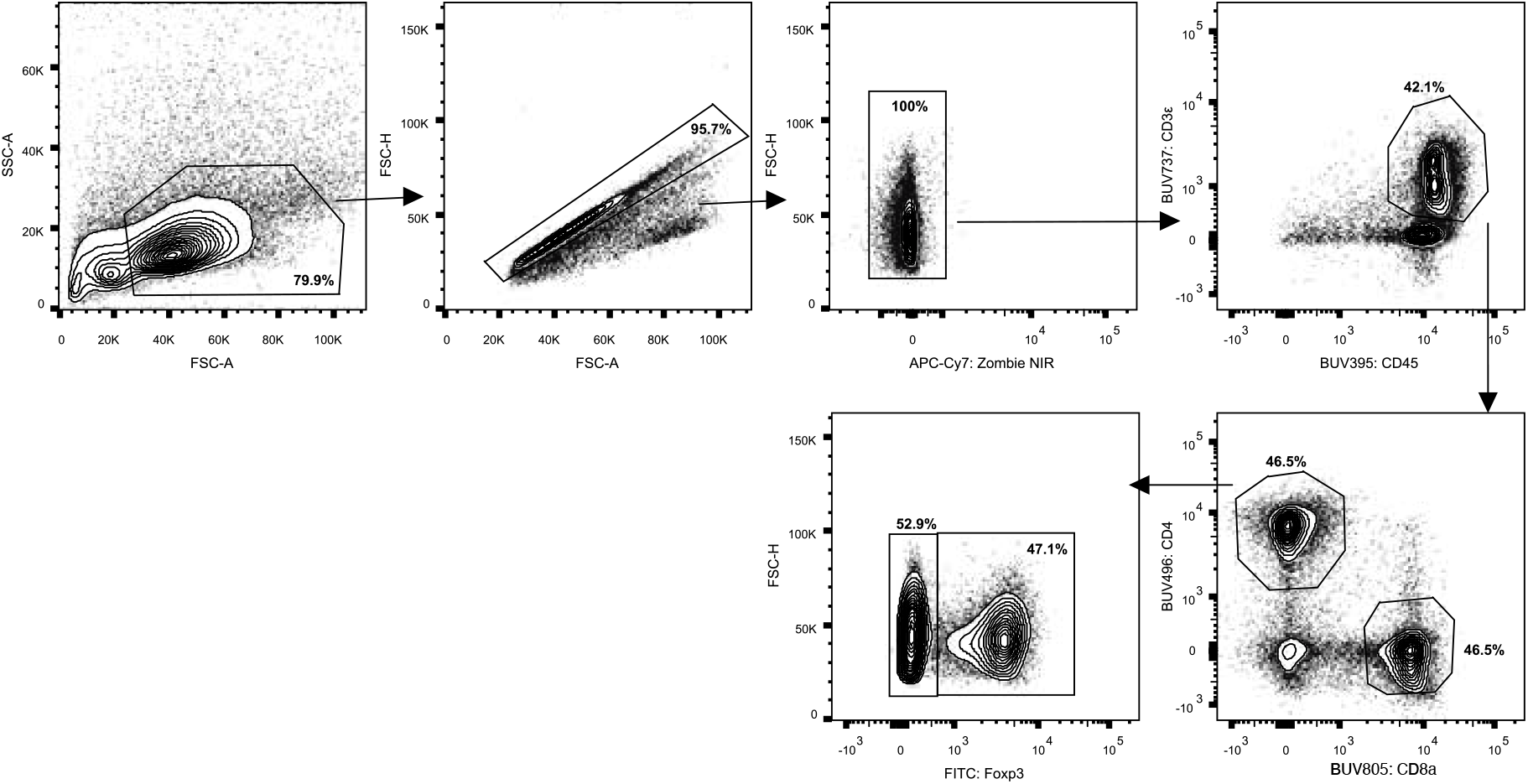
T cell identification flow cytometry gating strategy. Representative gating for CD8+ T cells, conventional T cells (Tcon; CD4+FOXP3-), and regulatory T cells (Treg; CD4+FOXP3+). All T cells are CD45+.

**Supplementary Figure 14.**
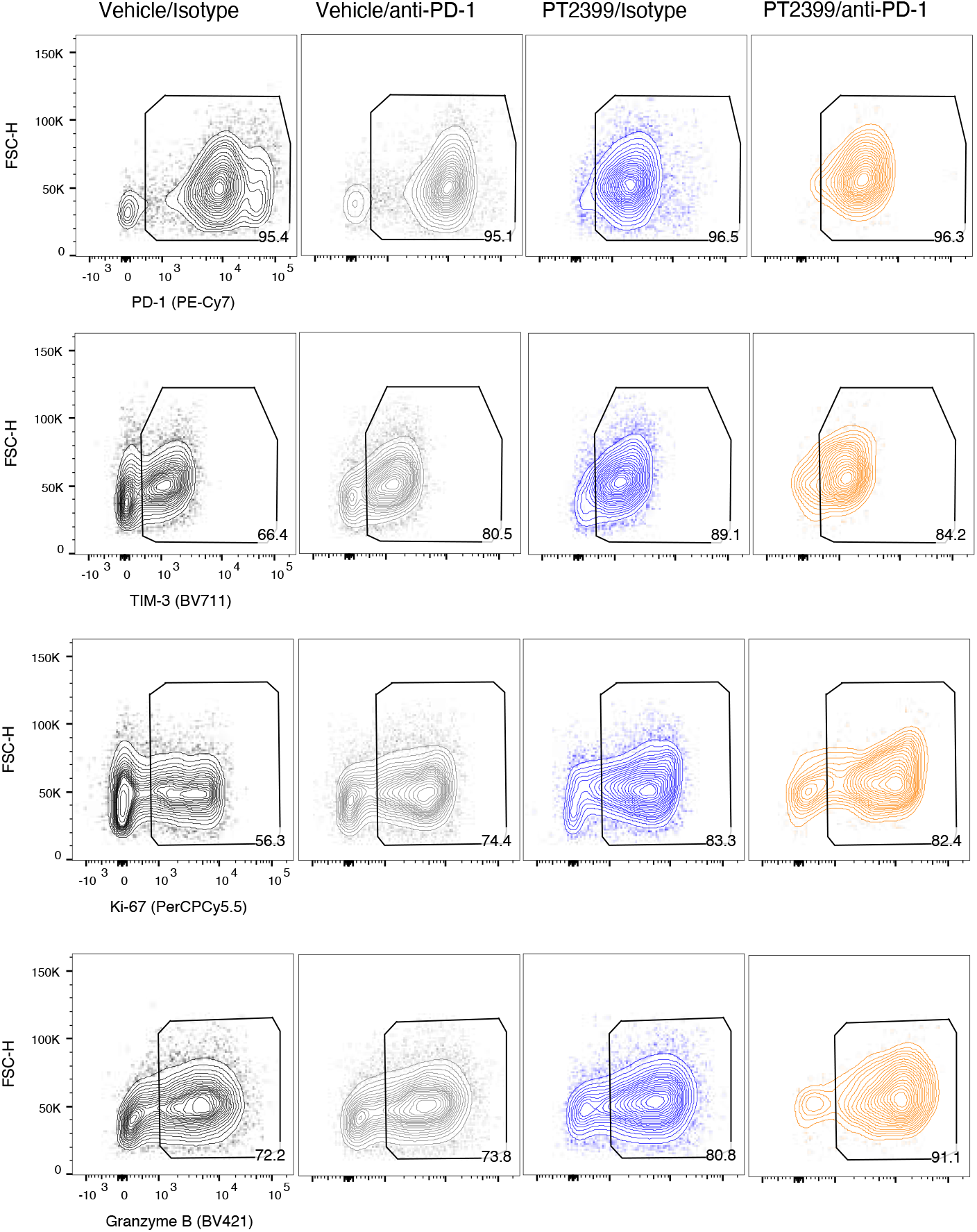
CD8+ cell characterization flow cytometry gating strategy. Representative gating for CD8+ T cell functional characteristics.

**Supplementary Figure 15.**
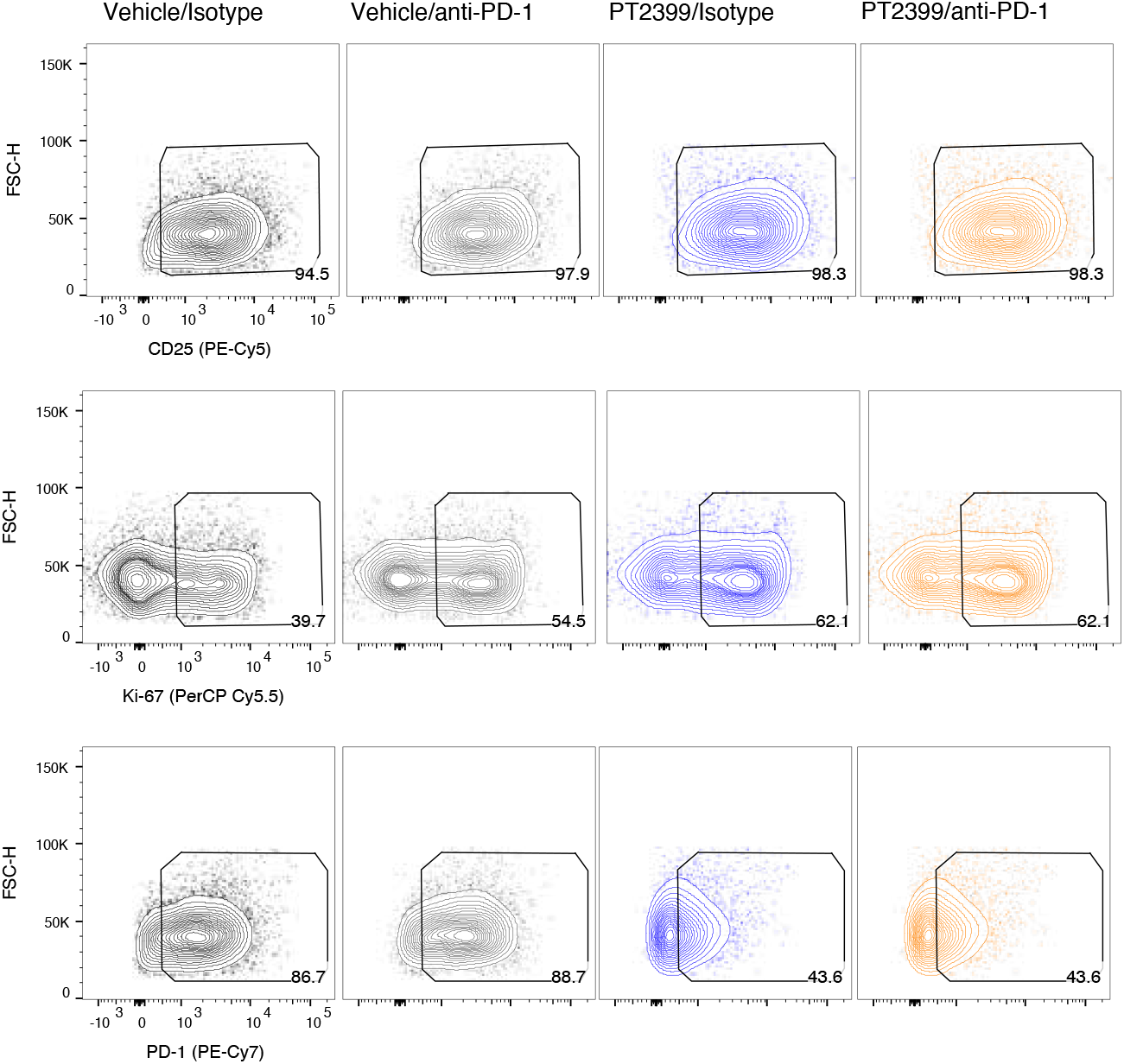
Regulatory T cell (Treg; CD4+FOXP3+) cell characterization flow cytometry gating strategy. Representative gating for regulatory T cell functional characteristics.

**Supplementary Figure 16.**
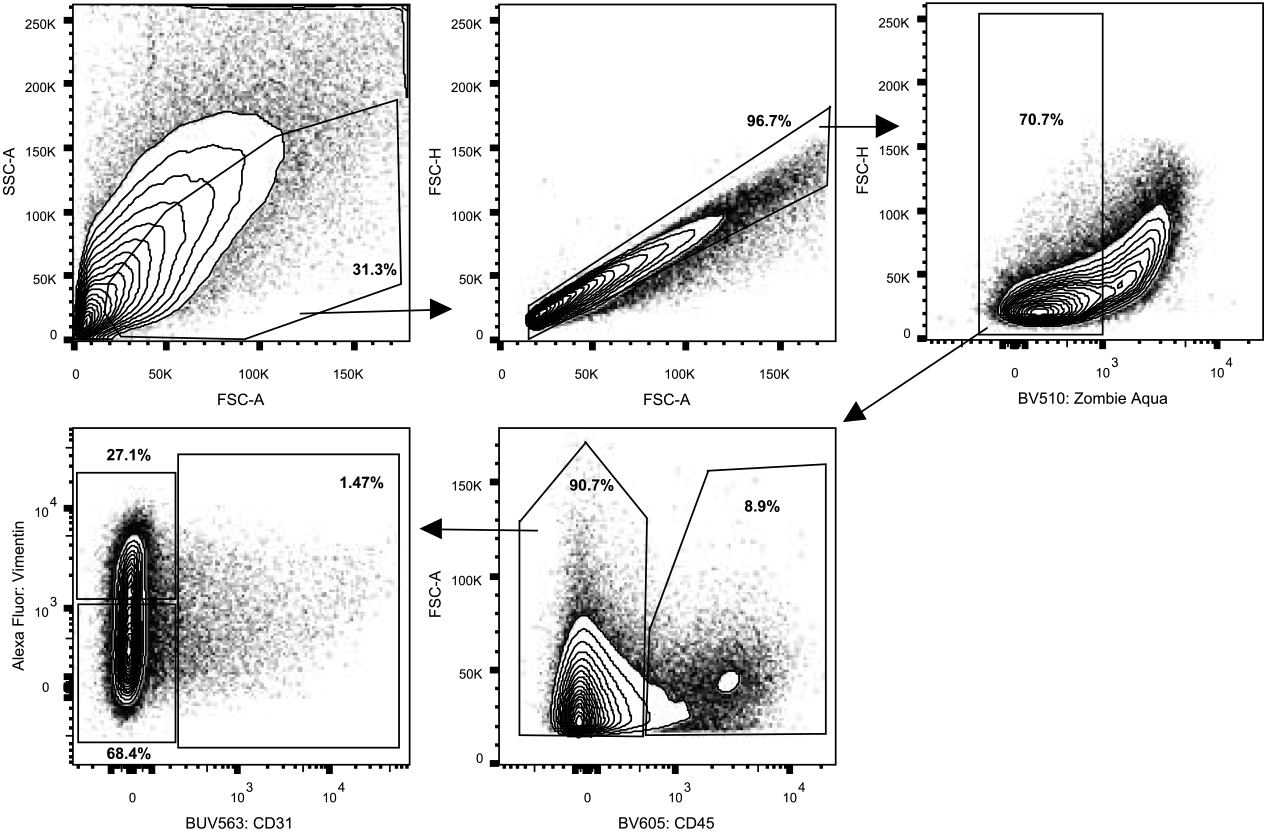
CD31+ cell identification flow cytometry gating strategy. Representative gating for CD45-CD31+ cells.

**Supplementary Figure 17.**
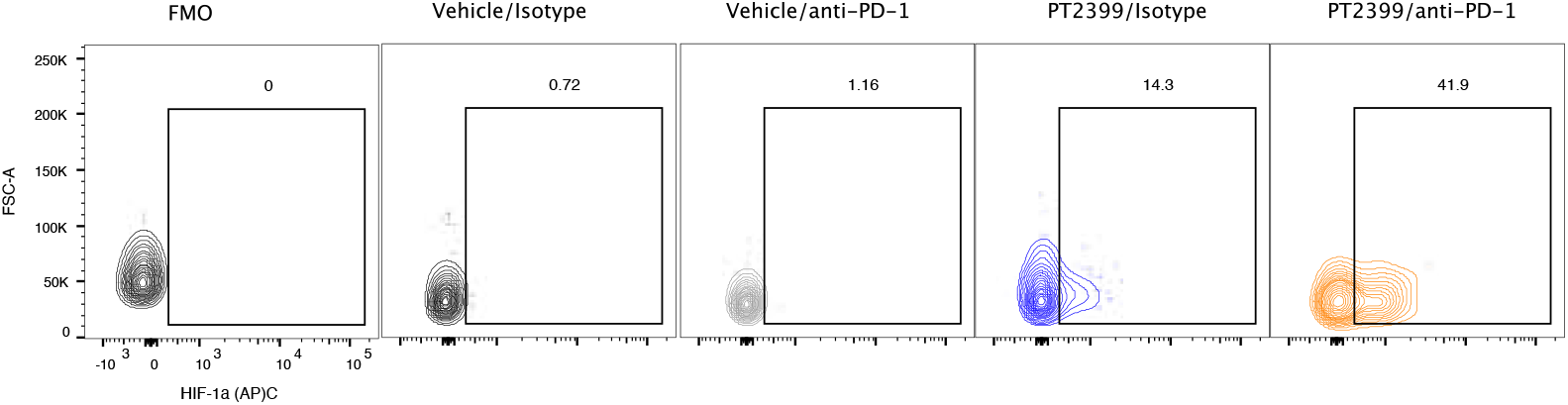
HIF-1*α*+CD31+ cell flow cytometry gating strategy. Example gating strategy of HIF-1*α*+ cells from total CD31+ cells.

